# Principles Governing the Dynamics of GABAergic Interneurons in the Barrel Cortex

**DOI:** 10.1101/554949

**Authors:** Jianing Yu, Hang Hu, Ariel Agmon, Karel Svoboda

## Abstract

Information processing in the neocortex is performed by GABAergic interneurons that are integrated with excitatory neurons into precisely structured circuits. To reveal how each neuron type shapes sensory representations, we measured spikes and membrane potential of specific types of neurons in the barrel cortex while mice performed an active, whisker-dependent object localization task. Whiskers were tracked with millisecond precision. Fast-spiking (FS) neurons were activated by touch with short latency and by whisking. FS neurons track thalamic input and provide feedforward inhibition. Somatostatin (SOM)-expressing neurons were also excited by touch, but with a delay (5 ms) compared to excitatory (E) and FS neurons. SOM neurons monitor local excitation and provide feedback inhibition. Vasoactive intestinal polypeptide (VIP)-expressing neurons were not driven by touch but elevated their spike rate during whisking, disinhibiting E and FS neurons. Our data reveal rules of recruitment for specific interneuron types, providing foundations for understanding cortical computations.

## INTRODUCTION

Modern molecular methods, in combination with neurophysiology and anatomy, have advanced our knowledge of the diversity in cortical GABAergic interneurons (Feldmeyer et al., 2018; Jiang et al., 2015; Rudy et al., 2011; Taniguchi et al., 2011; Tasic et al., 2018; Tremblay et al., 2016). But we are only beginning to unravel specific functions of different types of GABAergic interneurons during behaviors such as sensory processing, movement control, and cognition (Gentet et al., 2010, 2012; Kvitsiani et al., 2013; Lee et al., 2013; Muñoz et al., 2017; Sachidhanandam et al., 2016; Yu et al., 2016). We also need a much clearer picture of how interneuron types are recruited during sensory processing. To address these gaps, it is essential to measure the dynamics of defined types of interneurons on synaptic time-scales (e.g., milliseconds) during behavior.

In layers (L) 2-6 of the neocortex, parvalbumin (PV)-expressing FS interneurons, SOM interneurons, and VIP interneurons together represent about 82% of all GABAergic interneurons (Rudy et al., 2011). Different types of interneurons have unique laminar distributions (Feldmeyer et al., 2018). FS interneurons are concentrated in layers targeted by the ventral posteromedial nucleus (VPM), including L4, and the border that straddles deep L5 and upper L6 (L5b/6) (van Brederode et al., 1991). Other layers, including L2/3 and upper L5 (L5a), contain FS neurons at lower density. The density of SOM neurons increases with cortical depth, peaking in L5b/6 (Tremblay et al., 2016). VIP neurons are concentrated in L2/3 (Prönneke et al., 2015). The synaptic inputs and outputs of inhibitory neurons are cell type-specific (Figure 1A). In the barrel cortex, thalamocortical inputs from VPM or the posterior medial nucleus (POM) make strong connections with FS interneurons in L4/5b/6 or L5a, respectively, with very weak input to SOM interneurons (Agmon and Connors, 1991; Audette et al., 2018; Beierlein et al., 2003; Bruno and Simons, 2002; Cruikshank et al., 2007, 2010; Gabernet et al., 2005; Gibson et al., 1999; Porter et al., 2001; Swadlow, 1995). Locally, FS and SOM neurons both receive inputs from cortical excitatory (E) neurons (Beierlein et al., 2003). FS interneurons inhibit nearby FS, E and SOM neurons (Beierlein et al., 2003; Sun et al., 2006; Xu et al., 2013). SOM neurons inhibit both E and FS neurons (Beierlein et al., 2003; Ma et al., 2012; Xu et al., 2013). VIP neurons preferentially inhibit SOM neurons and are driven by long-range neuromodulatory and corticocortical inputs (Fu et al., 2014; Lee et al., 2013).

**Figure 1.**
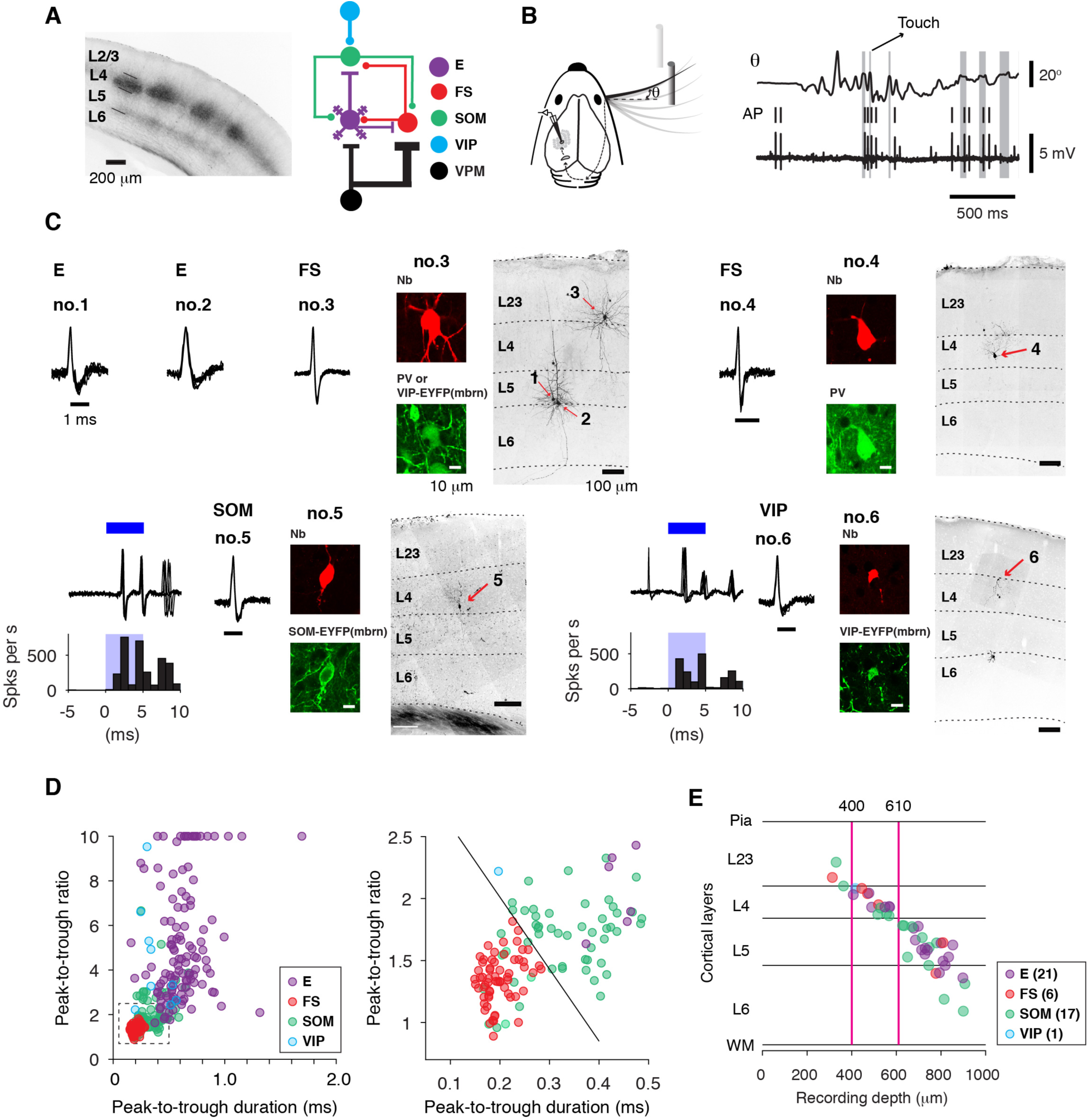
Cell Type-Specific Recordings During Active Tactile Behavior. (A) Left, coronal section through the barrel cortex with labeled VPM afferents (mCherry) in L4 and the border of L5 and L6. Right, connectivity of major interneuron classes with VPM inputs and local E neurons. (B) Left, object localization task. Mice move their whiskers to palpate a pole which is presented in one of two locations. Right, whisker position (*θ*) and action potentials (AP) recorded during the object localization task. Gray bars, periods of touch between whisker and pole. (C) Recordings from specific cell types. Cell types (excitatory, or E; fast-spiking, parvalbumin-expressing, or FS; somatostatin-expressing, or SOM; vasoactive intestinal polypeptide-expressing, or VIP) were identified based on a combination of spike waveform, immunohistochemistry, and optogenetic tagging in transgenic mice. Cells no. 1-3 were from VIP-IRES-Cre x Ai32 mice and the brain slices were immunostained against PV. In this section both VIP and PV cells emitted green fluorescence, but fluorescence in VIP cells was membrane-bound. Nb, neurobiotin. (D) Spike waveform parameters for different cell types. Peak-to-trough ratios larger than 10 are plotted as 10. (E) Laminar positions of neurons recovered histologically after recordings, and the corresponding recording depth (manipulator reading). The positions are normalized to the boundaries of the layers in which the cells were identified. Magenta lines denote 400 and 610 μm, corresponding to the boundaries for L4. The number of cells is listed for each type.

Transgenic mice expressing Cre recombinase in subsets of interneurons have provided a critical enabling tool for cell type-specific electrophysiology (Taniguchi et al., 2011). Studies using these mice have revealed the relationship between sensory stimuli or behavioral events and the activity of defined interneuron subtypes (Kvitsiani et al., 2013; Lee et al., 2013; Muñoz et al., 2017; Pi et al., 2013). In the somatosensory cortex, most previous recordings were targeted to GFP-expressing neurons by fluorescence microscopy and were therefore limited to superficial cortical layers (Gentet et al., 2010, 2012; Lee et al., 2013; Sachidhanandam et al., 2016). However, different cortical layers are innervated by different thalamic nuclei and long-range cortical inputs (Kinnischtzke et al., 2014; Lee et al., 2013; Lu and Lin, 1993; Mao et al., 2011; Petreanu et al., 2009; Wimmer et al., 2010), and have distinct local connectivity patterns (Bureau et al., 2006; Hooks et al., 2011; Jiang et al., 2015; Lefort et al., 2009; Shepherd and Svoboda, 2005). Discovering the principles governing the functions of specific types of interneurons, therefore, requires measurements across cortical layers (Muñoz et al., 2017). We employed a channelrhodopsin (ChR2)-guided optogenetic tagging strategy to home in on specific types of interneuron (Lima et al., 2009; Moore and Wehr, 2013; Muñoz et al., 2014; Yu et al., 2016), and recorded spikes or membrane potential of defined interneuron types across L2-6 during behavior.

To study active tactile processing, we trained head-restrained mice to use a subset of 1-3 whiskers to scan space for an object (pole) and report the result by licking (Guo et al., 2014a; Hires et al., 2015; O’Connor et al., 2010a, 2013; Yu et al., 2016). Inactivation experiments have shown that activity in the barrel cortex is used by the mice to perform the task (Guo et al., 2014a; Hong et al., 2018; O’Connor et al., 2010a, 2013). Using intrinsic signal imaging, we targeted our recordings to the cortical regions representing the whiskers used by the mice to perform the task. High-speed videography (Clack et al., 2012) allowed us to track whisker movement and touch with millisecond temporal precision and relate neural activity and behavior at these time-scales.

The whisker somatosensory system shows rapid activation to sensory input. VPM neurons respond to active touch with a 2-3 millisecond latency and with submillisecond jitter (Petersen et al., 2008; Yu et al., 2016) and relay this information to L4 and L5b/6 (Figure 1A). VPM neurons also increase their rate at the onset of whisking (Gutnisky et al., 2017; Moore et al., 2015; Poulet et al., 2012; Urbain et al., 2015; Yu et al., 2016). We tracked the propagation of these behavior-associated fast and slow signals from the VPM thalamus through different layers and cell types of the cortical circuit. Our measurements thus link the synaptic organization of a canonical cortical circuit with cell type-specific functions during active sensation.

### Highlights

- Recordings from major interneuron classes during active tactile behavior.
- Touch activates SOM neurons with a delay.
- Whisking, but not touch, excites VIP neurons.
- Whisking and touch activate FS neurons in the VPM thalamorecipient layers.

## RESULTS

### Cell-Type-Specific Recordings in Behaving Mice

We recorded from barrel cortex neurons while head-restrained mice performed a whisker-dependent object-localization task (Figure 1B). Whisker movements and the contacts between whisker and object were monitored using high-speed (1000 frames per s) videography (Clack et al., 2012; Hires et al., 2015; O’Connor et al., 2010a, 2013; Pammer et al., 2013; Yu et al., 2016). We performed single-cell juxtacellular recordings with glass pipettes, which provided unambiguous single-unit isolation (de Kock et al., 2007; O’Connor et al., 2010b; Yu et al., 2016) and the opportunity to label recorded neurons (Pinault, 1996). In some experiments, we performed membrane-potential recordings in the whole-cell mode with patch-clamp electrodes (Yu et al., 2016).

We recorded from E neurons and three inhibitory interneuron classes: FS, SOM, and VIP. E neurons (n = 153) had broad spike waveforms and a high peak-to-trough ratio (Fig. 1C, no 1, 2). We identified FS interneurons (n = 47) by their spike waveform (narrow peak-to-trough duration, low peak-to-trough ratio; Figure 1C, D). A subset of FS neurons (n = 3) was labeled by juxtacellular electroporation, and their identity was verified by immunohistochemistry against parvalbumin (Figure 1C, cell no. 3, 4). In our analysis we included 30 additional FS neurons from previous studies, recorded in VGAT-ChR2 mice (thin-spike ChR2+ cells, n =22), PV-IRES-Cre mice (PV+ cells, n = 4), PV-ChR2 mice (ChR2+ cells, n = 1), and C57BL/6 mice (n = 3)(Yu et al., 2016). To detect SOM (n = 80) or VIP (n = 9) neurons, we selectively expressed ChR2 in either type by crossing SOM-IRES-Cre or VIP-IRES-Cre mice (Taniguchi et al., 2011) with a ChR2 reporter line, Ai32 (Madisen et al., 2012), and exploited light-driven responses to identify these neurons (Figure 1C, cell no. 5, 6). A subset of SOM and VIP neurons were labeled to verify their recording locations (Figure 1C, E).

Spike waveform parameters of SOM neuron were intermediate between FS and E neurons, and overlapped with both populations, highlighting the importance of optogenetic tagging (Figure 1D). Additional information about the spike waveform allowed us to refine classification further, and exclude a small proportion (n = 6) of “pseudo SOM neurons” which were most likely FS neurons with off-target ChR2 expression (Figure S1) (Hu et al., 2013). VIP neurons had short spike durations but were distinguished from FS neurons by their high peak-to-trough ratios (Figure 1D). The subset of juxtacellularly labeled neurons (n = 45) was used to establish a correspondence between depth reading on our recording manipulator and laminar location of the recorded neurons (Figure 1E).

### Responses Evoked by Active Touch

We first analyzed responses to whisker touch. Contacts between whisker and the object elicited action potentials in E, FS, and SOM neurons in all layers (Figure 2). However, the response magnitudes (change in spike number within 50 ms after touch onset), response latencies (see Figure S2 for definition) and temporal dynamics differed across cell types and layers (Figure 3, Table 1). Touch-related input enters the barrel cortex via VPM, where neurons are driven by touch with short latencies (median, 2.6 ms; n = 29, Figure 3B, 3F) and low latency variability across cells (SD, 0.8 ms, n = 29, Figure 3B, I). In the VPM thalamocortical recipient layers of the barrel cortex (L4 and L5b/6), FS neurons responded with slightly longer latency (L4, 4.3 ms, n = 43; L5b/6, 5.6 ms, n = 15) and larger variability across cells (L4, 1.8 ms; L5b/6, 2.5 ms). E neurons responded with still longer latency (L4, 7.8 ms, n = 61; L5b/6, 8.7 ms, n = 26) and latency variability (L4, 5.1 ms; L5b/6, 5.4 ms). Stronger response magnitudes were seen in L5b/6 E (0.8 spikes per touch, n = 30) neurons compared to L4 E (0.1 spikes per touch, n = 95) neurons (*p* = 0.0054, Figure 3J) but the magnitudes were not different between L5b/6 FS (2.5 spikes per touch, n = 15) and L4 FS neurons (2.0 spikes per touch, n = 43; *p* = 0.21). In other layers, E and FS neurons responded with longer latencies than in L4 and L5b/6 (E: L2/3, 9.7 ms, n = 5, L5a, 10.3 ms, n = 16; FS: L2/3, 7.0 ms, n = 7, L5a, 7.1 ms, n = 7) and were thus likely excited by neurons in L4 and L5b/6 via interlaminar excitatory pathways or by a relayed pathway through the higher-order thalamus, POM (Groh et al., 2008). Touch-related VPM input, therefore, enters the cortex to excite FS and E neurons in L4 and L5b/6 simultaneously, consistent with previous reports with passive stimulation (Armstrong-James et al., 1992; Constantinople and Bruno, 2013; de Kock et al., 2007).

**Table 1.** Latency and magnitude of touch-evoked response.

|  |  | L2/3 | L4 | L5a | L5b/6 |
| --- | --- | --- | --- | --- | --- |
| E | Latency (ms) | 9.2 ± 1.2,<br>9.7, 8.1 – 10.1,<br>n = 5 | 9.4 ± 5.1,<br>7.8, 6.4 – 11.6,<br>n = 61 | 10.8 ± 3.3,<br>10.3, 8.7 – 13.2,<br>n = 16 | 10.6 ± 5.4,<br>8.7, 7.6 – 10.9,<br>n = 26 |
|  | Magnitude (spikes/touch) | 0.7 ± 0.6,<br>0.5, 0.3 – 0.8,<br>n = 5 | 0.2 ± 0.3,<br>0.1, 0.0 – 0.3,<br>n = 95 | 0.5 ± 0.5,<br>0.4, 0.1 – 0.9,<br>n = 23 | 0.8 ± 0.8,<br>0.8, 0.0 – 1.3,<br>n = 30 |
| FS | Latency (ms) | 6.7 ± 1.8,<br>7.0, 5.0 – 7.7,<br>n = 7 | 4.6 ± 1.8,<br>4.3, 3.6 – 4.8,<br>n = 43 | 7.5 ± 2.9,<br>7.1, 5.4 – 9.7,<br>n = 7 | 6.2 ± 2.5,<br>5.6, 4.2 – 8.0,<br>n = 15 |
|  | Magnitude (spikes/touch) | 2.4 ± 1.8,<br>2.9, 0.6 – 4.1,<br>n = 8 | 2.2 ± 1.0,<br>2.0, 1.5 – 3.1,<br>n = 43 | 1.8 ± 1.3,<br>1.2, 0.8 – 2.5,<br>n = 7 | 2.9 ± 1.6,<br>2.5, 1.7 – 3.6,<br>n = 15 |
| SOM | Latency (ms) | 13.9 ± 4.1,<br>12.5, 11.0 – 16.0,<br>n = 8 | 15.6 ± 4.1,<br>15.4, 12.4 – 17.2,<br>n = 18 | 13.2 ± 3.8,<br>12.7, 11.6 – 13.7,<br>n = 15 | 13.6 ± 3.6,<br>12.8, 11.5 – 14.3,<br>n = 17 |
|  | Magnitude (spikes/touch) | 0.5 ± 0.6,<br>0.5, 0.2 – 0.7,<br>n = 9 | 0.4 ± 0.4,<br>0.3, 0.1 – 0.6,<br>n = 27 | 0.5 ± 0.6,<br>0.3, 0.1 – 0.6,<br>n = 18 | 0.3 ± 0.4,<br>0.3, 0.0 – 0.6,<br>n = 26 |
| VIP | Magnitude (spikes/touch) | -0.2 ± 0.4,<br>-0.02, -0.4 – 0.1,<br>n = 9 |  |  |  |

- Values represent mean, median, 25<sup>th</sup>-75<sup>th</sup> percentiles, and cell number. - Response magnitude is defined as the average number of spikes after touch (0 – 50 ms), after subtracting the baseline spike rate (-25 – 0 ms from the touch onset).

**Figure 2.**
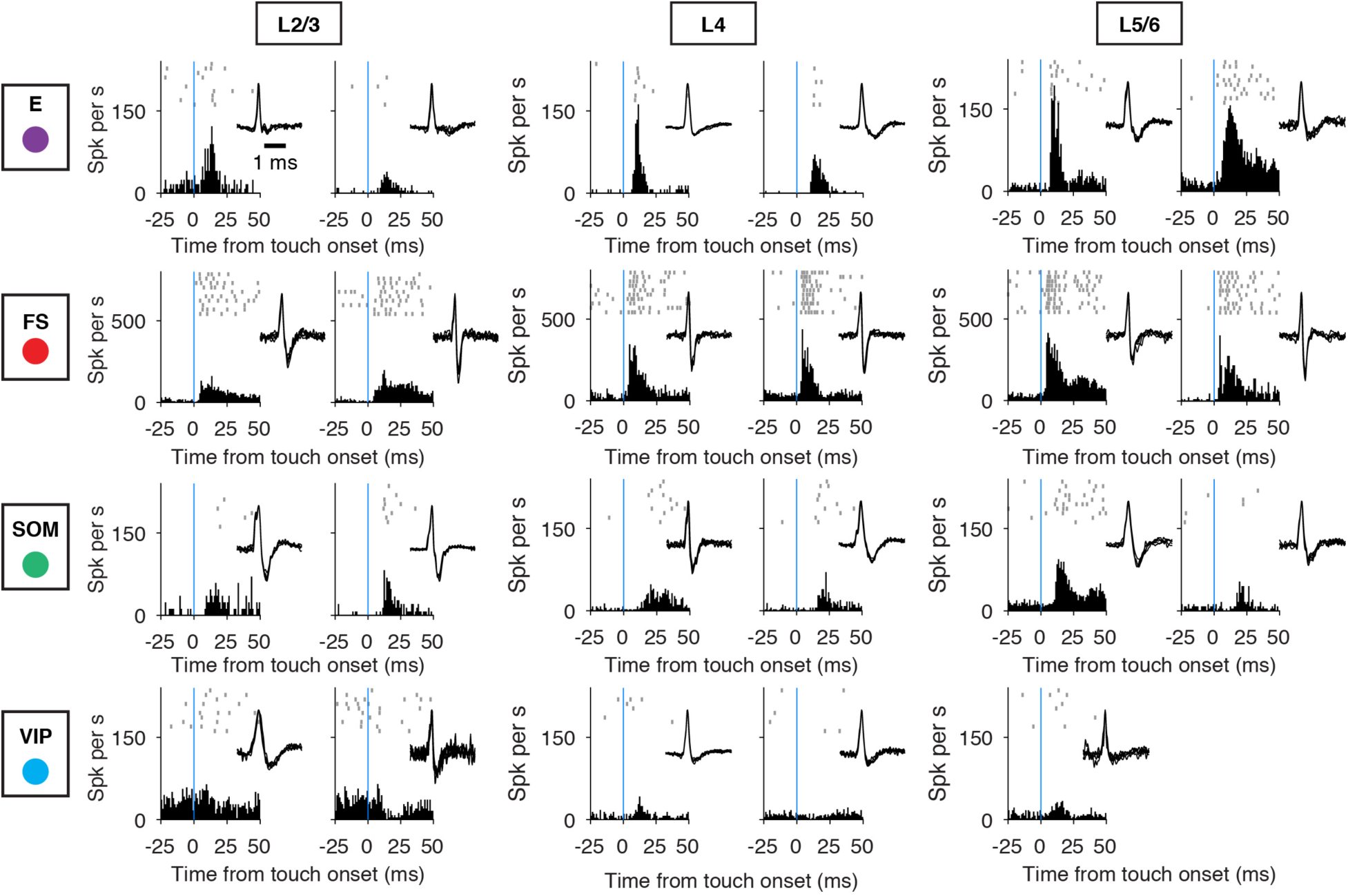
Cell-Type-Specific Activity Aligned to Touch Onset. Example recordings for different cell types (E, FS, SOM, VIP), arrayed vertically. Layers are arrayed horizontally. Each neuron is represented by spike raster plots across ten trials (top), the peristimulus time histogram (PSTH; bin size, 1 ms) aligned to touch onset (bottom), and the spike waveform (inset). Blue lines denote the touch onset.

**Figure 3.**
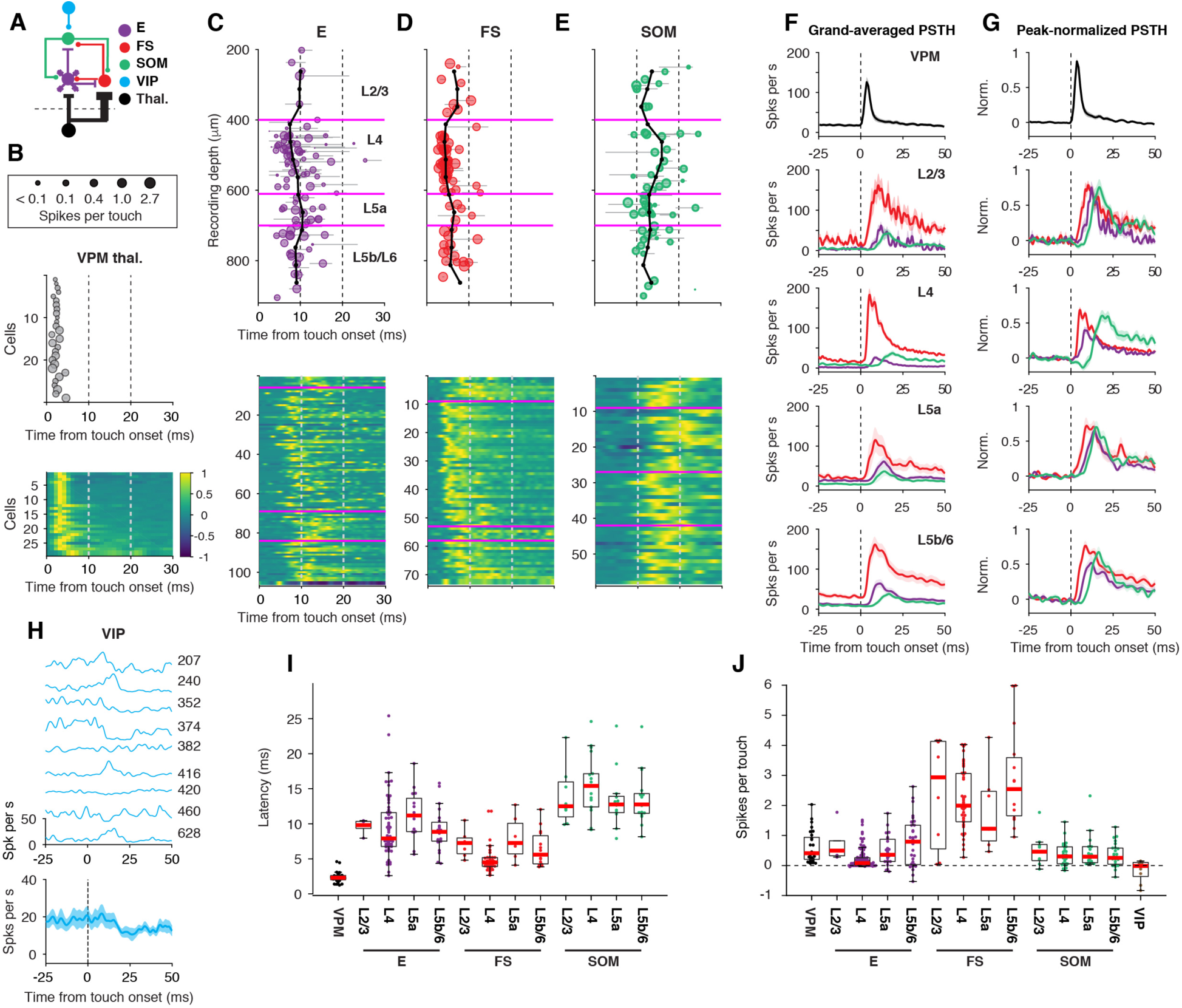
Touch-Evoked Spiking Responses Across Cell Types and Layers. (A) Connectivity diagram. (B) Response latency from touch onset for VPM neurons. Top, the size of each circle corresponds to the strength of the response (spikes per touch). Middle, touch latency. Bottom, peak-normalized PSTHs for all neurons. (C) Response latency from touch onset as a function of recording depth for excitatory (E) neurons. Top, the size of each circle corresponds to the strength of the response. Gray lines, 90% confidence interval for latency estimation for each cell. Magenta, 400, 610, and 700 μm representing the borders between L2/3 and L4, L4 and L5a, and L5a and L5b/6, respectively. Black lines, running median across cortical depth (calculated over 125 μm windows, in 50 μm steps). Bottom, peak-normalized PSTH for all neurons. (D) Same as C, fast-spiking (FS) neurons. (E) Same as C, somatostatin-expressing (SOM) neurons. (F) Spike rate aligned to touch onset for different neuron types (grand average; shading indicates SEM). (G) Same as F, peak-normalized. (H) Raw PSTHs for all VIP neurons and their grand average. Number is the recording depth (µm). (I) Response latency from touch onset for different neuron types and layers. VIP neurons did not respond to touch reliably and were not included. Boxplots follow the style of Tukey boxplot. (J) Response magnitude for all cell types. More cells, including those whose latency could not be determined, are included for this plot.

SOM neurons in all cortical layers were excited by touch, but with longer latencies than E or FS neurons (median, 13.2 ms, SD, 3.9 ms, n = 58, Figure 3E-G, I, Table 1). In L4, their responses were further delayed by early inhibition (median, 15.4 ms, SD, 4.1 ms, n = 18, Figure 3F, G). Touch elicited fewer spikes in SOM neurons compared to FS neurons across all cortical layers (Figure 3F, J, Table 1). In L4, for example, the response magnitude of SOM neurons (0.3 spikes per touch, n = 27) was several-fold smaller than for FS neurons (2.0 spikes per touch, p < 10^−7^) but was stronger than that of E neurons (0.1 spikes per s, p = 0.02).

Since most VPM neurons only fired briefly after touch (Figure 3B, F, G) (Yu et al., 2016), the long latency of excitation implies that SOM neurons are driven by cortical E neurons, and not directly by VPM input. The E → SOM neuron synapse is weak but facilitating (Beierlein et al., 2003; Gibson et al., 1999; Reyes et al., 1998). Facilitation by repetitive presynaptic spiking could occur as a result of repetitive touches. However, inter-touch-intervals (ITI) of successive touches were mostly too long for effective summation (median, 114 ms; the proportion of ITIs smaller than 25 ms, 2.1%; 82159 successive touches). Moreover, SOM neurons were fired effectively by the first touch of a whisking bout (Figure S3), making repetitive touches unnecessary for firing SOM neurons. Alternatively, it is possible that high-frequency (> 100 Hz) burst-like spiking, which was seen in a subset of E neurons (Figure S4), produces a rapidly facilitating depolarization in SOM neurons, triggering spikes. We tested this possibility in brain slice experiments (Figure S4). High-frequency bursting of E neurons, at frequencies (200-400 Hz) matched to *in vivo* recordings, elicited a rapid build-up of membrane depolarization in connected SOM neurons, reflecting both synaptic facilitation and postsynaptic charge summation (Figure S4). Additionally, the high connection probability of E→SOM (67% in our dataset of 36 tested L4 pairs) implies that simultaneous EPSPs from multiple presynaptic E cells will summate in single SOM neurons.

Because of their sparsity and their relatively small size (Prönneke et al., 2015), only a small number of VIP interneurons were recorded (L2/3; n = 5; L4, n =3; L5, n = 1). VIP interneurons were poorly driven by touch (median, −0.02 spikes per s, n = 9; Figure 2, Figure 3H, J, Table 1). The grand mean PSTH showed modest suppression by touch (Figure 3I, J), with a time-course and latency (18 ms) consistent with direct inhibition from SOM neurons (Pfeffer et al., 2013).

We next explored the synaptic mechanisms underlying touch-evoked spiking in E and SOM neurons using whole-cell recordings (Figure 4). Consistent with spike recordings, L4 E and FS neurons showed short-latency (median, L4 E, 4.4 ms, n = 37, L4 FS, 4.0 ms, n = 8) touch-evoked depolarization (Figure 4A-F) (Yu et al., 2016). L5b/6 E neurons show delayed depolarization overall (median, 7.0 ms, n = 31) but a subset was depolarized at short latency (~4 ms; e.g., cells 6 and 7 in Figure 4A), consistent with direct VPM input (Agmon and Connors, 1992; Constantinople and Bruno, 2013; Petreanu et al., 2009). L5/6 FS neurons showed responses that were virtually indistinguishable from L4 FS neurons, with rapid onset (median, 3.8 ms, n = 5).

**Figure 4.**
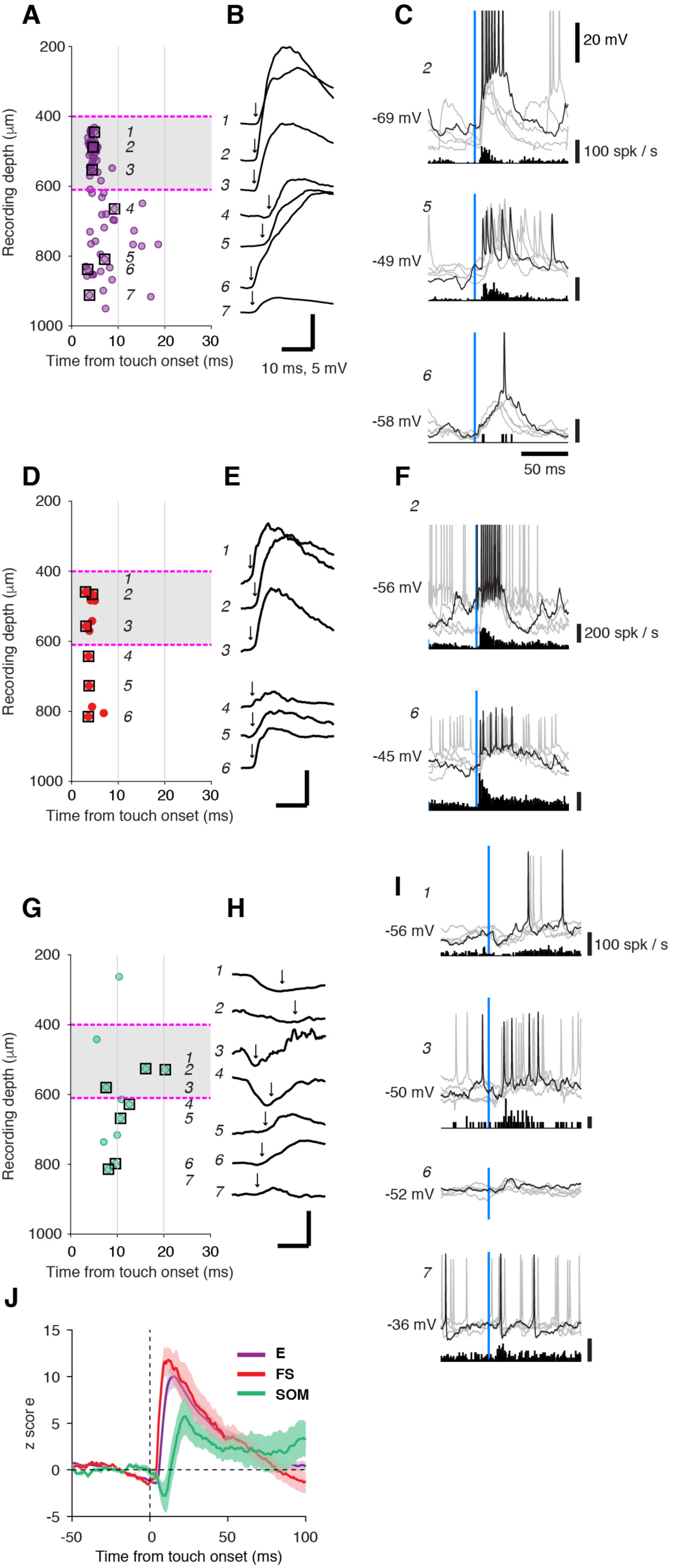
Touch-Evoked Synaptic Potentials Across Cell Types and Layers. (A) The latency of synaptic-potential depolarization after touch-onset as a function of recording depth, excitatory (E) neurons. Each point is an individual neuron. Squares correspond to example responses shown in B and C. (B) Example responses of 7 cells averaged across trials (seven cells, boxed in A). Arrow indicates the latency of synaptic-potential depolarization. (C) Single membrane potential traces and spike histogram for 3 cells. One of the trials is plotted in black and others in gray. (D-F) Same as (A-C) for FS neurons. (G-I) Same as (A-C) for SOM neurons. (J) Grand-averaged touch-evoked membrane potential response (z-scored) for E, FS, and SOM neurons.

Touch-evoked depolarization was delayed in SOM neurons (Figure 4J). Further inspection revealed two types of touch responses in SOM neurons. Responses of the first type (see cells 1-4 in Figure 4G and H) consisted of rapid hyperpolarization following touch, with an onset latency of 4 ms, coinciding with the onset of touch-evoked spiking in FS interneurons. Following this inhibition, delayed excitation drove these neurons to fire spikes (latency > 7 ms, Figure 4I, cells 1 and 3). These responses were found in L4 and L5a. Responses of the second type (see cells 5-7 in Figure 4G and H), found in L5/6, consisted of a delayed excitatory input (latency, 8-10 ms), with no evidence of early inhibition. For both types, the delayed depolarization suggested that SOM neurons were driven by cortical E neurons.

### Whisker Movement-Dependent Responses

Active whisker touch requires voluntary movement of the whiskers. Whilst touch activates sensory receptors in the periphery, movement can modulate activity in the sensory system through centrally-generated signals related to movement commands or arousal (Fu et al., 2014; Lee et al., 2013; Poulet et al., 2012), in addition to activation of peripheral receptors, or reafference (Campagner et al., 2016; Severson et al., 2017; Yu et al., 2016). These diverse signals can modulate specific types of interneurons and shape the state of the cortical network prior to touch. We analyzed the modulation by whisker movement of activity in all four cell types across cortical layers by aligning activity to the onset of bouts of whisking (Figures 5, 6).

**Figure 5.**
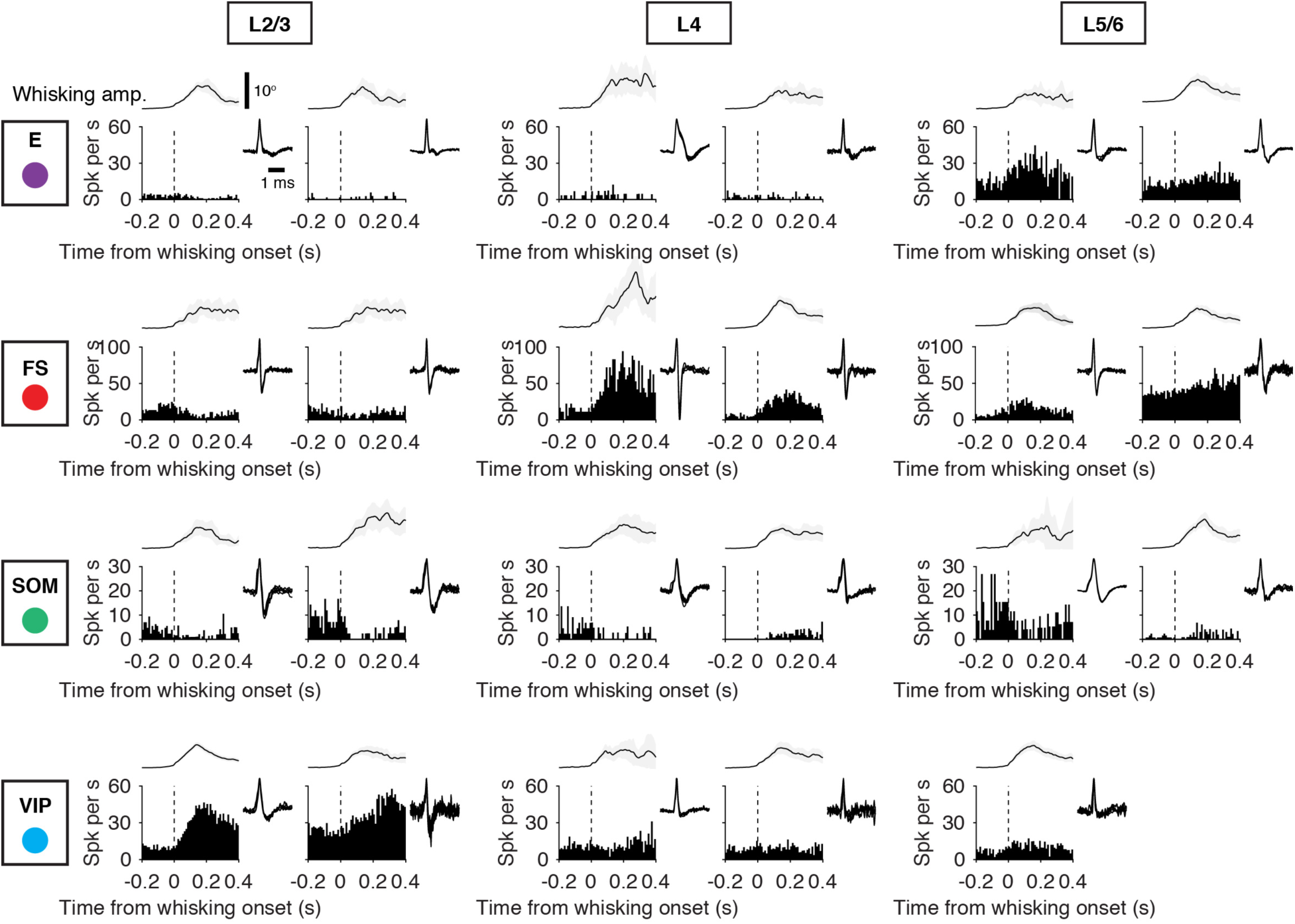
Cell Type-Specific Activity Aligned to Onset of Whisker Movement. Example recordings for different cell types (E, FS, SOM, VIP), arrayed vertically. Layers are arrayed horizontally. Each neuron is represented by the PSTH aligned to the onset of a whisking bout and the spike waveform (inset). Top, whisking amplitude.

**Figure 6.**
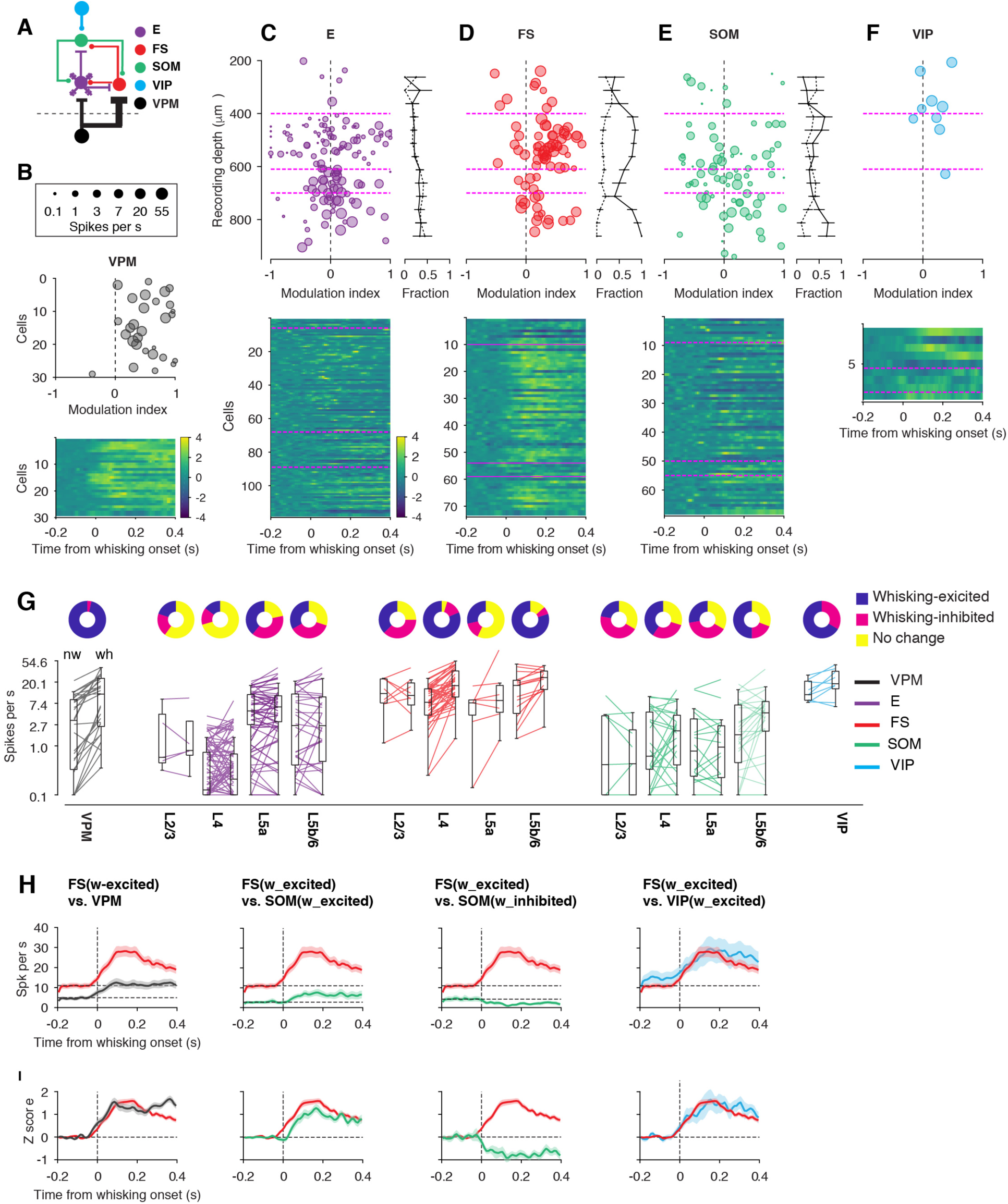
Whisking-Evoked Responses Across Cell Types and Layers. (A) Connectivity diagram. (B) Top, modulation index for individual VPM neurons. The size of the marker corresponds to the spike rate, averaged between whisking and non-whisking periods. Bottom, averaged responses (z-scored) for individual neurons aligned to the start of whisking bouts. (C) Top left, modulation index for E neurons as a function of recording depth. The size of the marker corresponds to the spike rate. Right, the proportion of neurons excited by whisking (solid line) and those inhibited by whisking (dashed line) calculated in running windows (125 μm window, 50 μm step). Error bar denotes the stand error. Bottom, averaged responses (z-scored) for individual neurons aligned to the start of whisking bouts. (D) Same as C for FS neurons. (E) Same as C for SOM neurons. (F) Same as C for VIP neurons. (G) Changes in spike rate with whisker movements for individual cells, arranged by cell types and layers. Pie charts, the fraction of cells that is excited (blue), inhibited (red), or shows no change in spike rate (yellow) to whisking. (H) Spike rate changes aligned to start of whisking bouts for VPM (n = 29), FS (L4 and L5b/6, whisking-excited, n = 47), SOM (L4 and L5b/6, whisking-excited, n = 24, whisking-inhibited, n = 13), and VIP neurons (n = 9), plotted pairwise for ease of comparison. Cells excited by whisking are plotted separately from cells inhibited by whisking. (I) Same as H with z-scored spike rate.

VPM neurons are strongly excited by whisking (Figure 6B) (Yu et al., 2016). Barrel cortex E neurons were excited or inhibited, in roughly equal proportions throughout the cortical layers (Figure 6C, G). On average, the spike rate of E neurons did not change with whisking (Table 2). As reported previously (Yu et al., 2016), a majority (81.4%) of L4 FS interneurons were driven by whisker movement through VPM input (Figure 6D). In L5b/6, where VPM afferents also terminate (cf. Figure 1A), a similar proportion (80.0%) of FS neurons showed increased spike rate to whisker movement. FS neurons in other layers (L5a, L2/3) did not show consistent responses to whisking. A significant fraction of L2/3 FS neurons (3/8, or 37.5%) was inhibited by whisking (Figure 5, L2/3, FS) (See also Gentet et al., 2010).

**Table 2.** Spike rates during non-whisking and whisking periods.

|  |  | L2/3 | L4 | L5a | L5b/6 |
| --- | --- | --- | --- | --- | --- |
| E | Non-whisking (spikes/s) | 2.7 ± 3.7,<br>0.6, 0.5 – 4.5,<br>n = 5 | 0.5 ± 0.8,<br>0.1, 0.0 – 0.7,<br>n = 95 | 6.8 ± 5.2,<br>5.2, 2.7 – 11.2,<br>n = 23 | 6.1 ± 6.9,<br>2.6, 0.4 – 11.5,<br>n = 30 |
|  | Whisking (spikes/s) | 2.5 ± 3.9,<br>0.8, 0.7 – 3.2,<br>n = 5 | 0.7 ± 1.2,<br>0.2, 0.0 – 0.7,<br>n = 95 | 7.7 ± 6.2,<br>6.3, 3.1 – 10.6,<br>n = 23 | 7.8 ± 10.6,<br>2.6, 0.5 – 10.0,<br>n = 30 |
| FS | Non-whisking (spikes/s) | 13.8 ± 8.9,<br>11.7, 7.5 – 23.3,<br>n = 8 | 10.2 ± 7.2,<br>7.8, 4.3 – 14.7,<br>n = 43 | 7.5 ± 5.2,<br>7.6, 4.3 – 8.7,<br>n = 7 | 16.9 ± 14.3,<br>17.2, 4.6 – 22.0,<br>n = 15 |
|  | Whisking (spikes/s) | 14.4 ± 11.9,<br>10.7, 6.8 – 19.3,<br>n = 8 | 21.3 ± 14.6,<br>17.0, 10.0 – 34.0,<br>n = 43 | 20.6 ± 32.7,<br>8.5, 4.9 – 17.2,<br>n = 7 | 24.3 ± 11.7,<br>24.9, 14.2 – 33.8,<br>n = 15 |
| SOM | Non-whisking (spikes/s) | 2.6 ± 3.6,<br>0.4, 0.03 – 4.1,<br>n = 9 | 2.6 ± 3.2,<br>0.6, 0.3 – 4.9,<br>n = 27 | 2.8 ± 4.5,<br>0.8, 0.2 – 3.6,<br>n = 18 | 3.9 ± 4.9,<br>1.7, 0.5 – 6.9,<br>n = 26 |
|  | Whisking (spikes/s) | 1.3 ± 1.7,<br>0.4, 0.03 – 2.1,<br>n = 9 | 3.0 ± 3.4,<br>2.0, 0.4 – 4.2,<br>n = 27 | 2.8 ± 5.4,<br>0.9, 0.2 – 2.6,<br>n = 18 | 6.1 ± 7.5,<br>4.7, 2.0 – 8.1,<br>n = 26 |
| VIP | Non-whisking (spikes/s) | 14.6 ± 7.3,<br>11.1, 8.5 – 21.0,<br>n = 9 |  |  |  |
|  | Whisking (spikes/s) | 23.0 ± 13.2,<br>18.6, 14.7 – 33.4,<br>n = 9 |  |  |  |

- Values represent mean, median, 25<sup>th</sup>-75<sup>th</sup> percentiles, and cell number.

SOM interneurons exhibited diverse responses to whisking (Figure 5 and 6E). In L2/3, whisking reduced the spike rate of a majority of SOM neurons (44.4%) and increased the rate of a smaller subset (22.2%). In L4 and L5a, SOM neurons were inhibited (L4, 29.6%, L5a, 38.9%) or excited (L4, 40.7%, L5a, 50.0%) by whisking. In L5b/6, a majority of SOM neurons (50.0%) showed a spike rate increase and a minority showed a decrease (19.2%). Following whisking onset, compared to VPM (latency to half-height of the peak, 27 ms) and FS (40 ms) neurons, the spike rate increases of SOM neurons in the thalamorecipient layers (L4 and L5b/6) were further delayed (57 ms) (Figure 5, Figure 6H, I). This slow response likely reflects cholinergic input acting through muscarinic acetylcholine receptors (Chen et al., 2015; Muñoz et al., 2017; Xu et al., 2013).

A majority of VIP neurons (66.7%) also elevated their spike rate during whisking (Figure 6F, G). The lack of touch-driven activity in VIP neurons argues that whisking drives these neurons by cholinergic input acting on nicotinic receptors (Fu et al., 2014) or long-range excitatory input from the motor cortex (Lee et al., 2013). In spite of different underlying mechanisms, VIP and FS neurons increased their firing rate with similar dynamics during whisking (Figure 6H, I).

### Cell-Type-Specific Disinhibition by VIP Interneuron

VIP neurons mainly target SOM neurons, and to a lesser extent, FS neurons (Hioki et al., 2013; Lee et al., 2013; Pfeffer et al., 2013). It is difficult to predict how the activation of VIP neuron, for example, during whisking, will influence the activity of FS and E neurons. Although activation of VIP interneuron can produce disinhibition in the cortical circuits (Lee et al., 2013; Pi et al., 2013), this has not been shown directly in the barrel cortex *in vivo* and the laminar specificity is unknown. Thus, we optogenetically stimulated VIP neurons while measuring the activity of FS, E, and putative SOM neurons.

We expressed ChR2 in VIP interneurons and applied high-frequency (5 ms pulses at 50 Hz) photostimuli to increase spike rate of VIP neurons. Long (3 seconds) photostimuli produced tonic inhibition of putative SOM neurons (Figure S5, 7C). VIP neuron stimulation activated L2/3 (100%), L4 (70%) and L5/6 (50%) FS neurons, in particular at the beginning of stimulation (0-170 ms from the laser onset). Photostimulation of VIP interneurons increased the spike rate of L5/6 E neurons preferentially, with little effect on L2/3 and L4 E neurons (Figure 7A, B and C). These results suggest that VIP neurons activation during whisking will produce a cell-type-and layer-specific effect at a slow time-scale (100 ms).

**Figure 7.**
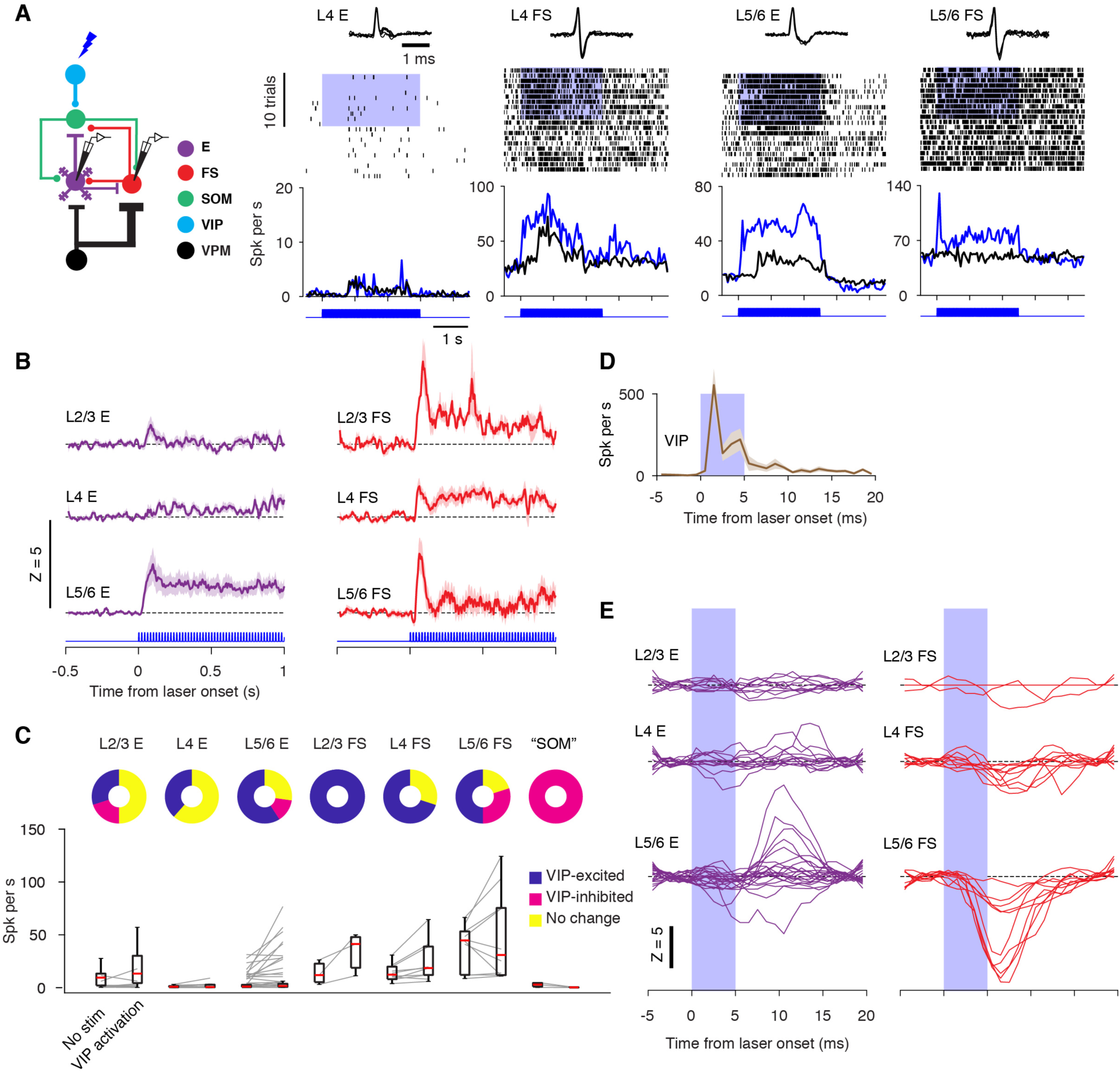
Activation of VIP Neurons Causes Cell Type-Specific Disinhibition and Inhibition. (A) Examples of responses of E and FS neurons to photostimulation of VIP neurons. Top, spike waveforms. Middle, spike rasters. Blue shading, time of photostimulation. Bottom, spike rate under control condition (black) and with VIP photostimulation (blue). Activity is aligned to the beginning of trials and the modulation under the control condition (black tick marks and traces) was caused by the behavior (e.g., touch). (B) Grand-averages of the spike rate changes (stim-control, z-scored) of E (L2/3, n = 10; L4, n = 13; L5/6, n = 22) and FS (L2/3, n = 3; L4, n = 10; L5/6, n = 10) neurons in response to VIP neuron stimulation, sorted by their layers. (C) Effects of VIP photostimulation on neurons in different layers. (D) Responses of VIP neurons to single laser pulses (n = 9). (E) Responses of E and FS neurons to VIP stimulation, aligned to single laser pulses.

Single laser pulses triggered a brief increase of spike rate (2-5 ms latency after photostimulus onset) in VIP neurons (Figure 7D, 1C, cell no. 6). We analyzed the spike rate modulation in E and FS neurons locked to individual laser pulses to uncover direct inhibition from VIP neurons. Each laser pulse reduced the spike rate of L5/6 FS interneurons, revealing selective monosynaptic inhibition of L5/6 FS interneurons (z score at 5.5 ms, −7.8 ± 1.6, mean ± SD, *p* = 0.002, n = 10; Figure 7E). L4 FS neurons were also inhibited by VIP neurons directly, but at a relatively smaller magnitude (−2.0 ± 0.6, *p* = 0.014, n = 10). In contrast, individual E neurons in all layers showed virtually no direct inhibitory response to single VIP pulses (−0.5 ± 0.21, *p* = 0.75, n = 45), demonstrating a lack of direct inhibition from VIP to E neurons. The strong direct inhibition from VIP interneurons is likely the cause that, in a subset of L5/6 FS cells (30%), the firing rate was suppressed by prolonged VIP activation. Presumably, in this subset, the direct inhibition overpowered the indirect disinhibitory effect via the VIP→SOM→FS pathway, an effect that dominated and caused increased FS firing in other layers.

## DISCUSSION

We discovered principles governing the functional recruitment of major subtypes of GABAergic inhibitory interneurons in the somatosensory cortex. The activity of FS neurons resembled that of thalamic input, whereas SOM neuron activity followed the local excitatory neuron population. VIP neurons were not driven by sensory signals but still were modulated during behavior by non-sensory pathways.

### FS Interneurons Track the Thalamic Input

FS neurons in the VPM projection layers (L4 and L5b/6) were activated by touch at short latencies and were sensitive to whisker movement (Figure 6), matching the dynamics of the VPM input (Yu et al., 2016). FS interneurons thus have similar functions across VPM thalamorecipient layers: filtering bottom-up inputs from the VPM thalamus, suppressing movement-associated reafferent activity, and narrowing the “window of opportunity” for excitatory neurons to respond to touch (Cruikshank et al., 2007; Gabernet et al., 2005; Simons and Carvell, 1989; Yu et al., 2016). FS neurons in L5a, innervated by POM afferents (Audette et al., 2018; Lu and Lin, 1993), were not consistently driven by whisker movement, which may reflect the weak representation of whisker movement in POM (Masri et al., 2008; Moore et al., 2015; Urbain et al., 2015). In L2/3, FS interneurons showed delayed activation after touch, consistent with their activation by ascending input from L4 or L5 (Dantzker and Callaway, 2000).

We demonstrated that VIP neuron activation, through inhibiting SOM neurons, excited FS neurons across all layers, including L4 (Fig. 7). Since the activation of VIP neurons is likely independent of sensory input, the VIP-SOM-FS pathway provides a means by which centrally-generated signals influence the firing rate of FS interneurons in the sensory cortex.

### SOM interneurons integrate inputs from local circuits

Touch elicited spiking responses in SOM neurons throughout L2-6, in spite of the fact that they lack appreciable thalamic innervation (Audette et al., 2018; Beierlein et al., 2003; Cruikshank et al., 2010). Touch-evoked response in SOM neurons was weaker than in FS neurons (Figure 3J) and showed delayed onset latency (Figure 3I). These features suggest that SOM neurons monitor the activity of local excitatory neurons, providing feedback inhibition once local excitation rises. Touch-evoked responses in SOM neurons could thus shape the duration of spiking in E and FS neurons.

A previous report, using two-photon-guided patching, only found touch-evoked suppression of L2/3 SOM neurons (Gentet et al., 2012). The discrepancy between our and their work is likely attributed to layer-or subtype-specific properties of SOM neurons. Previous work used GIN mice for two-photon-targeted recordings and recorded exclusively the upper portion of L2/3 (Gentet et al., 2012). In GIN mice, only a subset of SOM neurons in the barrel cortex express GFP (Ma et al., 2006). Here in our experiments using SOM-IRES-Cre mice (Taniguchi et al., 2011), touch-evoked spikes were found in SOM neurons across layers 2-6. In addition, our dataset does not include the upper portion of L2/3 (< 200 µm), which is likely where the majority of cells were collected by Gentet and colleagues.

SOM neurons are also subject to modulation by other pathways, such as cholinergic excitation (Chen et al., 2015; Muñoz et al., 2017; Xu et al., 2013) and inhibitory inputs from various types of interneurons, including FS and VIP neurons (Ma et al., 2012; Muñoz et al., 2017; Pfeffer et al., 2013). The spike rate of SOM neurons is thus set by the combined effects of these different input sources. In contrast to a previous report (Muñoz et al., 2017), which reports that L4 SOM neurons increase their spike rate by whisking, we found a substantial fraction (30%) of L4 SOM neurons was inhibited by whisking. This difference may stem from the difference in the behavioral state of animals: In our task, animals were constantly engaged in an active tactile task, where the FS neuron network might be more active during whisking, exerting a stronger inhibition on SOM neurons.

### VIP interneurons modulate the spike rate of cortical circuits

VIP interneurons were highly active spontaneously *in vivo* during behavior, even in the absence of touch and whisking (Fig. 6G, Table 2). Whisking further increased the spike rate of a majority of VIP interneurons. Despite their high baseline spiker rate, which implies that the membrane potential of VIP interneurons hovered around the spike threshold, touch did not strongly drive these neurons. Some VIP neurons were actually suppressed by touch with a latency that matched the activation of SOM neurons (Fig. 3H). VIP neurons are known to receive inhibitory inputs from both FS and SOM neurons (Pfeffer et al., 2013; Sohn et al., 2016; Staiger et al., 1997). Therefore, despite the fact that VIP interneurons receive excitatory inputs from the thalamus (Lee et al., 2010; Staiger et al., 1996) and local E neurons (Sohn et al., 2016; Staiger et al., 2002), the effective reversal potential caused by touch, which is a combination of both excitatory and inhibitory inputs (Crochet et al., 2011), is likely close to the baseline membrane potential of VIP neurons or even more hyperpolarized. Therefore, active touch is unable to increase the spike rate of VIP neurons further. Under other conditions, for example, passive whisker deflection, touch seems to increase the firing rate of VIP neurons in the upper portion of L2/3 (Sachidhanandam et al., 2016).

VIP neuron activation disinhibited E and FS neurons in the barrel cortex, via inhibition of SOM neurons (Figures 7, S5) (Jackson et al., 2016; Lee et al., 2013; Pi et al., 2013). Thus, the high spike rate of VIP neurons plays a role in constantly suppressing SOM neurons and reducing inhibition in the circuits, with or without whisking. We observed a strong laminar dependence that had not been reported before: E neurons in L5/6, not L2-4, increased their spike rate strongly in response to VIP neuron stimulation. The majority of FS neurons in all layers were disinhibited by VIP neuron activation, but a direction inhibition from VIP to FS neurons was more prevalent for L5/6 FS neurons (Dávid et al., 2007; Hioki et al., 2013), and was likely the reason that a sizeable fraction of L5/6 FS neurons decreased their firing rate upon VIP stimulation

### Conclusion and Outlook

We have established key principles underlying the recruitment of GABAergic inhibition during active behavior, revealing functional specificity of each interneuron type. Our study highlights the importance of cell-type-specific electrophysiology as a critical tool for unraveling functional relationships between neural activity and behavior at fine time-scales, which cannot be substituted with other methods, including monosynaptic tracing (Wall et al., 2016), or calcium imaging (Dipoppa et al., 2018; Pakan et al., 2016). Our data constrain parameters for theoretical and computational models of cortical circuits and pave the way for future investigation of the cortical circuits underlying various tactile functions and dysfunctions.

## METHODS

### Animals

All procedures were in accordance with protocols approved by the Janelia Research Campus Institutional Animal Care and Use Committee. We crossed SOM-IRES-Cre (JAX #013044) and Ai32 (JAX #012569) mice for targeted recording of SOM neurons and crossed VIP-IRES-Cre (JAX #010908) and Ai32 mice for targeted recording of VIP neurons. We also included data from a previous publication (Yu et al., 2016), in which these mouse strains were used for targeted recordings of GABAergic neurons: VGAT-ChR2-EYFP (JAX #014548), PV-IRES-Cre (JAX #017320), and PV-ChR2 (JAX #012355).

### Behavioral training

Mice (2-6 months old of both sexes) were implanted with a custom-made titanium head bar (11524, Flintbox). After 2-7 days of recovery, they were placed under water-or food-restriction schedule. Method of water restriction has been described previously (Guo et al., 2014b). For food restriction (DOI: dx.doi.org/10.17504/protocols.io.x9nfr5e), mice were provided with 3 g of food pellets (3.43 kcal/g, PicoLab Rodent Diet 20 5053) daily and were allowed to drink water *ad lib*. Weight was maintained at above 85% of *ad lib* fed mice. During behavioral training, Similac Advance Infant Formula (0.64 kcal/ml) was used as the reward, delivered through a peristaltic pump (P720, Instech Laboratories, Inc., PA), at a rate of 7 µl per reward. Since mice can consume up to 4 ml of infant formula but only 1 ml of water during a behavioral session, using infant formula prolonged the length of a behavioral session by more than 2 folds (>1200 trials per session). After the behavioral session, mice were supplemented with food pellets to meet their total daily calorie need of 10 kcal.

Mice were trained on a whisker-based go/nogo task (O’Connor et al., 2010a; Yu et al., 2016). A 0.5-mm diameter pole (class ZZ gage pin, Vermont Gage) was presented in an anterior location or one of several posterior locations. Mice were trained to lick an electrical lickport to receive rewards when the pole was in one of the posterior locations and withhold licking when the pole was in the anterior position. Trials were randomized and no more than 3 trials of the same types were presented in succession. All but 1-3 whiskers of C or D row were trimmed to facilitate whisker tracking. Recordings were targeted in the whisker somatosensory cortical regions of the spared whiskers, determined by transcranial intrinsic signal imaging before the recording sessions.

Whiskers were tracked with a Basler A504k camera using a telecentric lens (Edmund Optics) and 940-nm illumination (Roithner Laser). For each trial, 5 seconds of whisker movement were captured at 1000 fps with 100-150 µs exposure time, using Streampix 3 or 6 software (Norpix). Whisker trajectories and contacts between the whiskers and the pole were quantified using the Janelia Whisker Tracker (Clack et al., 2012) and custom-made scripts in Matlab (MathWorks).

### Neural recordings

#### *In vivo* juxtacellular recording

Detailed protocols (DOI: dx.doi.org/10.17504/protocols.io.wwkffcw) can be found at www.protocols.io (See also Yu et al., 2016). On the day of recording, small (200 µm-diameter) craniotomy was made in the targeted cortical region following a published method (Pinault, 2005). The dura was left intact in most cases. After surgery, the craniotomy was covered with a small drop of 1.5% agarose dissolved in a “cortex buffer” solution (in mM, NaCl 125, KCl 5, dextrose 10, HEPES 10, CaCl_2_ 2, MgSO_4_ 2, 272 mOsm, pH 7.4), followed by Kwik Cast (WPI) filling the entire recording chamber. Animals were allowed to recover on a warm blanket (37°C) for at least two hours before the recording session started.

For juxtacellular recording, long-shank borosilicate pipettes (1.50 mm OD/0.86 mm ID, Sutter Instruments, CA) were filled with electrolytes such as Ringer’s solution (in mM, NaCl 135, KCl 5.4, HEPES 5, CaCl_2_ 1.8, MgCl_2_ 1, 294 mOsm, pH 7.3) or 0.5 M NaCl, with 2% Neurobiotin (Vector Laboratories). The cortex was penetrated with a normal angle. The manipulator was zeroed when the pipette tip touched the surface of the brain, which often resulted in an increase in pipette resistance. To search for spikes from single cells, the appearance of spontaneous spikes or an increase in pipette resistance was monitored. To target ChR2-expressing neurons brief (5 ms) laser pulses (peak power, 2-5 mW) were delivered at low frequency (5 Hz, 3 seconds out of every 10 seconds). Laser-evoked spikes were monitored with an oscilloscope, triggered on the laser onset. ChR2-expressing neurons usually fired 1-2 ms after laser onset and were readily identifiable. During the searching process, drops of rewards were delivered at a rate of 7 µl per minute to keep the animals alert. The behavioral program was initiated once the desired neuron was found. The neural signal was amplified (Multiclamp 700B, Molecular Devices) and digitized at 10 kHz or 20 kHz, controlled by Ephus (Suter et al., 2010) or Wavesurfer (http://wavesurfer.janelia.org).

After the recording was complete, to fill the recorded cells, 1-10 nA positive current (50% duty cycle, 2 Hz) was injected to entrain the spikes of the recorded cell (Pinault, 1996). The evoked firing and recording noise were closely monitored. The current amplitude was adjusted online to avoid damage to the cell. After 5-15 minutes of juxtacellular filling, the animal was killed within 2 hours on the same day, or within 24 hours on the second day. Not more than one cell was labeled within 200 μm in the dorsal-ventral axis and 70 μm in the anterior-posterior or medial-lateral axis.

#### *In vivo* whole-cell recording

Whole-cell patch pipettes (6-9 MΩ) were filled with an internal solution (in mM, K-gluconate 135, KCl 4, HEPES 10, EGTA 0.5, Na_2_-phosphocreatine 10, Mg-ATP 4, and Na2-GTP 0.4, 292-300 mOsm, pH 7.3; or K-gluconate 125, KCl 7, HEPES 10, EGTA 0.5, Na_2_-phosphocreatine 10, Mg-ATP 4, and Na_2_-GTP 0.4, 292-300 mOsm, pH 7.3) with 0.4% Neurobiotin. The signal was amplified (Multiclamp 700B, Molecular Devices) and digitized at 10 kHz or 20 kHz, controlled by Ephus (Suter et al., 2010) or Wavesurfer (http://wavesurfer.janelia.org). For ChR2-guided whole-cell recording, the presence of laser-evoked spikes was examined in every instance when the pipette tip contacted a cell. If there were laser-evoked spikes, gigaseal and whole-cell configuration were established. Otherwise, the ChR2-cells were cleared with the ‘Buzz’ function (100 μs) in Multiclamp 700B.

#### *Ex-vivo* paired whole-cell recordings

Coronal brain slices of the somatosensory (S1) cortex were prepared as previously described (Ma et al., 2012), from 2.5-4 weeks old mice of both sexes. Most animals were of the X94 transgenic line (Ma et al., 2006) (JAX #006334) in which about half of all L4-5 SOM interneurons express GFP. Some experiments used progeny of the SOM-Ires-Cre line (JAX #013044) and the Ai9 tdTomato reporter (JAX #007909) (Madisen et al., 2010). Dual whole-cell recordings were performed from pairs of neighboring L4 excitatory neurons and SOM interneurons; SOM interneurons were targeted by their fluorescence, excitatory neurons by their soma shape, and the identity of both cell types was verified post-hoc by their characteristic firing patterns and membrane parameters (Hu and Agmon, 2016). High-frequency (200 and 400 Hz) spike trains were triggered in the excitatory neuron by brief (1 ms) current steps and trains of evoked unitary EPSPs (uEPSPs) were recorded from the postsynaptic interneuron, from a holding potential of about −70 mV (uncorrected for pipette junction potential). A total of 36 pairs were tested, 24 of which (67%) were synaptically connected in the E→SOM direction, including 15 (42%) connected bidirectionally. For analysis, we selected 10 pairs with a E→SOM connection tested at both 200 and 400 Hz and with at least one uEPSP >0.5 mV. Measurements were done on averages of (typically) 10-15 trials repeated at 8 s intervals.

### Optogenetic stimulation

The recording site (craniotomy) was photostimulated with a 473-nm laser (CL-473-150, Crystal Laser). The laser was gated by a shutter (Vincent Associates) and an acousto-optical modulator (AOM; MTS110-A3-VIS, Quanta Tech) and was focused on the craniotomy (beam diameter, 1.4 mm 99.9% energy or 4 × s.d.). Light pulses (5 ms) were delivered at either 5 Hz (for optogenetic tagging) or 50 Hz (for VIP neurons stimulation) with a maximal power of 5 mW.

### Analysis

#### Spike sorting and waveform

Spike waveforms were identified by amplitude thresholding, resampled to 40 kHz, and aligned by their peak. Putative waveforms were then subjected to principal component analysis (PCA) analysis and non-spike events (e.g., those associated with licking artifacts) were deleted. Putative spikes were inspected again, and additional non-spike events were deleted. To control for drift, only spikes whose peak-to-trough duration fell between 25^th^ to 75^th^ percentiles of all spikes from that cell were included for spike waveform analysis. These spike waveforms were then averaged for each cell and peak-normalized. The peak-to-trough duration (Figure 1) was defined as the time from the first peak to the first trough after the peak. The peak-to-trough ratio was defined as the ratio of the absolute values of peak and trough.

To examine the spike waveforms further (Figure S1), additional spike parameters were analyzed. Spike rise time was defined as the time from 25% of the peak, to the peak. Rising slope modulation was defined as 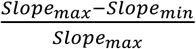. These two parameters separated quasi-FS SOM neurons from PV/FS neurons that expressed Cre transiently during development in SOM-IRES-Cre mice (Pseudo-SOM)(Hu et al., 2013). Pseudo-SOM neurons showed functional properties (e.g., response latency to touch) similar to FS neurons.

#### Whisking

The whisker angle was band-pass filtered between 6-60 Hz with a Butterworth filter (4th order) and decomposed using the Hilbert transformation. Phase ± π corresponds to the most retracted whisker location and phase 0 corresponds to the most protracted whisker location. Whisking epochs were defined as whisking cycles when the whisking amplitudes at phase 0 exceeded 2.5°. Non-whisking periods were defined as periods of at least 100 ms with negligible whisker movement (amplitude <1.25°), in the absence of touch or licking. Average spike rates during non-whisking (*SR_non-whisking_*) and whisking (*SR_whisking_*) periods were estimated from extracted non-whisking or whisking epochs. For individual cells, the statistical significance in whisking-induced modulation was evaluated by the Wilcoxon rank sum test (Figure 6G, pie chart). Whisking-induced modulation of spike rate was quantified by modulation index (Figure 6B-F):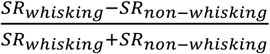. Transitions from non-whisking to whisking (e.g., Fig. 5 and 6) were defined as the end of a non-whisking epoch followed by a period in which the average whisking amplitude increased by least 2° and no touch occurred within 250 ms. If touch occurred after 250 ms, the data after touch were removed (replaced by NaN).

Spike rate was aligned to the whisking onset to build peri-stimulus time histograms using 10-ms bins (PSTH) (Figure 5). To normalize whisking-aligned PSTH, the baseline spike rate (from −200 to −50 ms before whisking onset) was subtracted from the PSTH. The baseline-subtracted PSTH was then normalized by the standard derivation of the entire PSTH (−200 to 400 ms from whisking onset). The z-scored PSTH was then smoothed with a Gaussian kernel (width, 1 bin).

#### Touch response, amplitude, and latency

The onsets and offsets of touches were curated manually by custom-made Matlab scripts. Touches were not included if their inter-touch-intervals (time between the offset of the previous touch and the onset of the current touch) were less than 25 ms. Passive touches, when the pole directly struck a resting whisker, were not included. Otherwise, all touches, including protraction-and retraction-induced touches, were included for analysis. Spike rate was aligned to the touch onset to build a PSTH using 1-ms bins (PSTH) (Figure 5). The touch-evoked response magnitude was defined as the average spike number between 0-50 ms after touch onset, with expected spike number from baseline activity subtracted (2 × average spike number between −25 – 0 ms relative to the touch onset).

To track the latency of spike increase after touch, spike times were convolved with a Gaussian kernel (1-ms width) to derive a spike density function (SDF, Figure S2), which were then averaged across trials (Nawrot et al., 1999). The peak response was defined as the maximal spike rate after touch onset (0-50 ms). Response latency was defined as the first time when the averaged SDF crossed the half-height of the peak response. If the peak response did not exceed 2.5 × SD of the baseline activity (estimated from - 25 ms to touch onset), the cell was excluded from latency analysis. To control reliability, bootstrap analysis (2000 resampling) was carried out. A 90% confidence interval was calculated from the bootstrap estimates for each cell and was plotted in Figure 3C-E. Cells were excluded for latency analysis if the 90% confidence interval of their latency estimation was larger than 15 ms.

Membrane-potential latency (Figure 4) was defined as the time from touch onset to the time when dVm/dt crossed the baseline (3 × SD of the dVm/dt between 0 and 25 ms before touch). For SOM neurons, since a sharp touch-evoked depolarization was not observed, the latency was manually determined as the time when Vm depolarized after the touch onset.

### Histology

Detailed protocols (DOI: dx.doi.org/10.17504/protocols.io.wa4fagw) on biocytin/neurobiotin staining and immunohistochemical staining of neurochemical markers such as parvalbumin can be found on procotol.io. Following the final recording session, mice were deeply anesthetized with isoflurane and perfused with 0.1 M phosphate buffer followed by 4% paraformaldehyde (PFA, in 0.1 M phosphate buffer, pH 7.4). The brain was immersed in fixative for at least 24 hours before sectioning. Cortical slices (100 µm) were collected from the areas surrounding the recording sites. To reveal biocytin/neurobiotin, brain slices were reacted overnight with 1:100 Streptavidin, Alexa Fluor 594 conjugate (1 mg/ml, S11227, Thermo Fisher Scientific) in a PBS solution containing 2% Triton X-100 (X100-1L, Sigma Aldrich), 20% DMSO (BP231-1, Fisher Scientific), and 4% Bovine Serum Albumin (B4287-5G, Sigma Aldrich). To detect neurochemical markers, immunohistochemical staining was performed with the primary antibody against parvalbumin (mouse, PV235, Swant) for 4-7 days at room temperature in above PBS solution, followed by secondary antibody staining using Alexa Fluor conjugates (Thermo Fisher Scientific) at room temperature for 4-7 days. Slices were air-dried and mounted on slides with Vectashield (H-1200 or H-1000, Vector Laboratories). DAPI or Neurotrace 435/455 (Thermo Fisher Scientific) was included for nucleus staining when applicable. Cells were examined with a confocal microscope, using 20x or 60x objectives (Zeiss 880). Laminar boundaries were determined by inspecting DAPI, NeuN, PV, or auto-fluorescent signals.

### Statistical analysis

Statistical comparisons were performed using the Wilcoxon rank sum test (unpaired). All statistical analyses were performed in Matlab (MathWorks).

## ACKNOWLEDGMENTS

This work was funded by the Howard Hughes Medical Institute (to K.S.). VPM data were collected by Diego Gutnisky and were published (Gutnisky et al., 2017; Yu et al., 2016). We thank Tina Pluntke for assistance in developing food restriction protocol and animal training. We thank Lupeng Wang for his advice on food restriction.

## Supplemental Information

Principles Governing the Dynamics of GABAergic Interneurons in the Barrel Cortex

Jianing Yu, Hang Hu, Ariel Agmon and Karel Svoboda

**Supplemental Figure 1.**
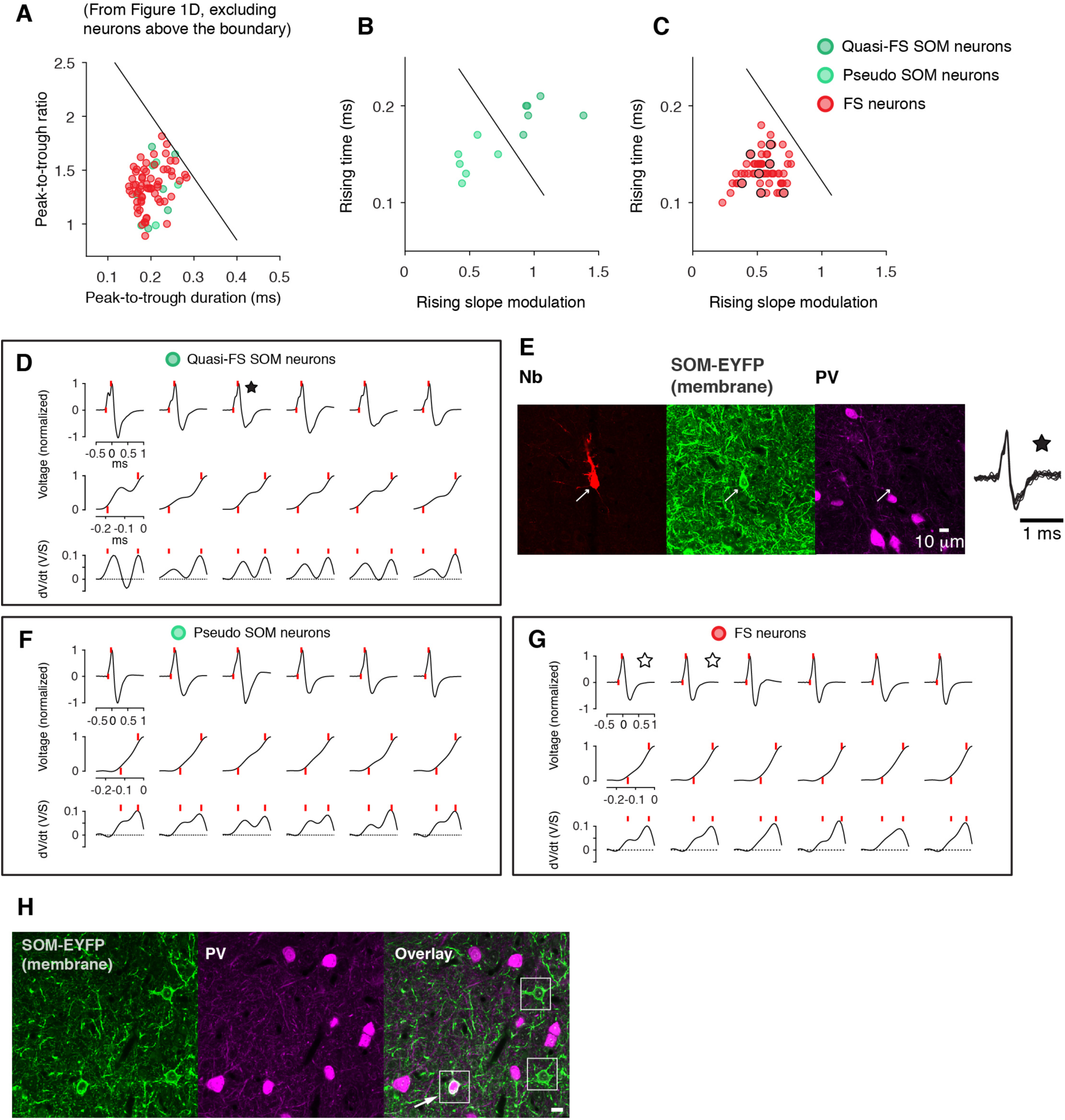
Identification of Thin-Spike SOM Interneurons. A subset of recorded ChR2+ neurons in SOM-IRES-Cre x Ai32 mice had thin spike waveform (e.g., they overlapped with FS neurons in terms of peak-to-trough duration and peak-to-trough ratio). Some of these neurons are likely quasi-FS SOM neurons. These neurons have been described in X-94 mice, in which non-Martinotti SOM neurons are labeled (Ma et al., 2006). Others could be FS/PV neurons that expressed ChR2 in an off-target manner (Hu et al., 2013). We distinguished these two possibilities based on two additional parameters extracted from the spike waveforms. (A) Thin-spike ChR2+ neurons recorded in SOM-IRES-Cre x Ai32 mice and FS neurons. (B) Thin-spike ChR2+ neurons were divided into two groups with two further parameters extracted from the spike waveform. The light green neurons symbols correspond to FS neurons with the off-target expression of ChR2 (pseudo SOM neurons), whereas the dark green symbols correspond to quasi-FS SOM neuron. (C) Same analysis as B for FS neurons. Circles mark those whose spike waveforms are shown in G. (D) Quasi-FS SOM neuron. The rising phase of these neurons always showed a brief decrease in the slope, or a “shoulder”. (E) One quasi-FS SOM neuron (marked with a star in D) was negative for PV immunoreactivity. Note the characteristic rising slope that contains a “shoulder”. (F) Putative FS neurons with an off-targeted expression of ChR2 (“pseudo SOM cells”) were distinguished by their rapidly rising slope that lacks a “shoulder”, similar to FS interneurons (G). (G) Spike waveforms of FS interneurons. Star-marked neurons were labeled and were test positive for PV immunoreactivity. (H) As previously reported, in SOM-IRES-Cre x Ai32 transgenic mice, some PV+ neurons express ChR2 (arrow), possibly owing to a transient Cre recombinase expression during development.

**Supplemental Figure 2.**
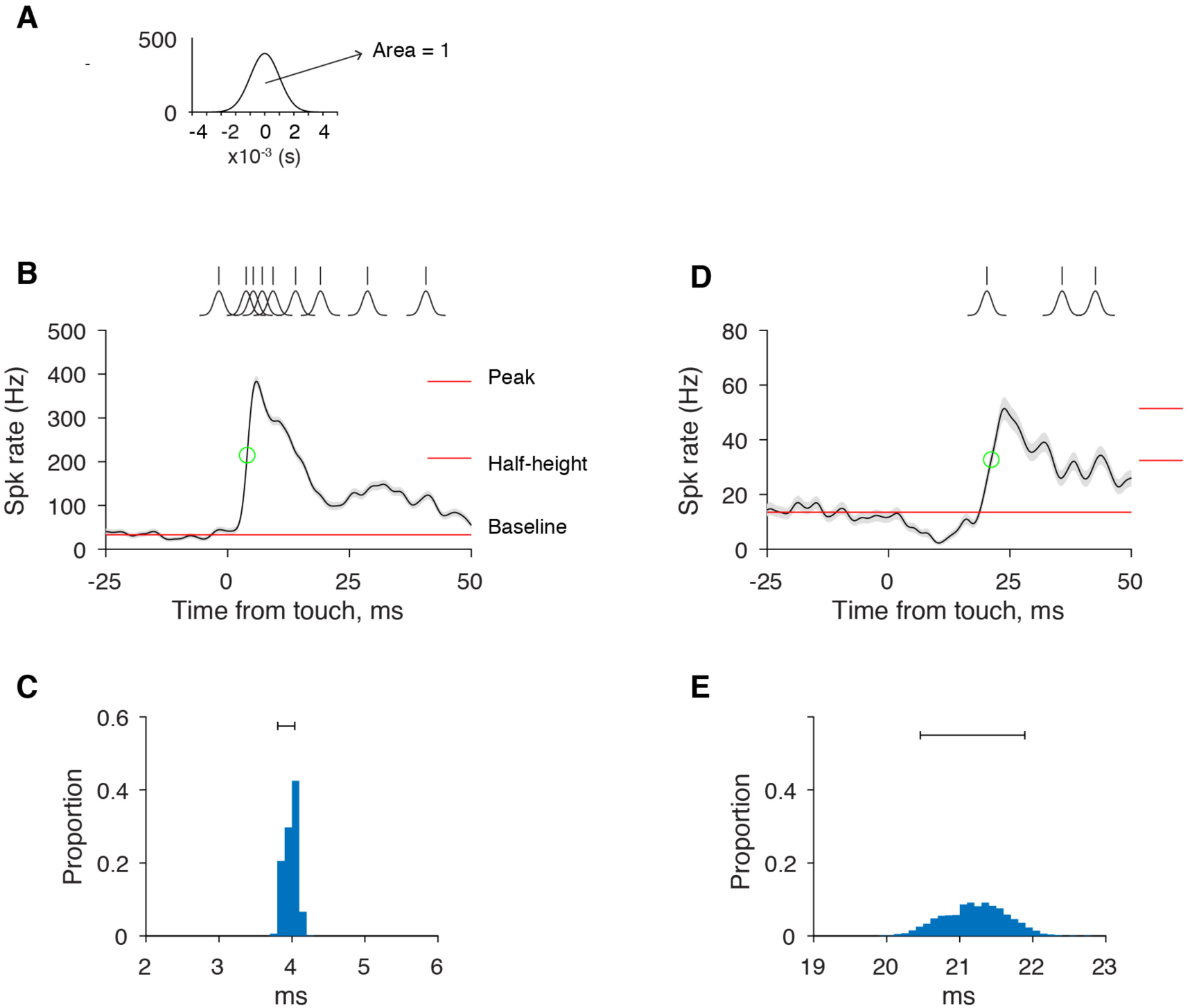
Latency of Touch Evoked Responses. (A) Profile of a Gaussian kernel (width=1 ms). The area under the curve is 1. (B) Spike trains of a FS neuron were convolved with the Gaussian kernel and averaged across trials to produce a spike density function. For latency estimation, the baseline, peak response, and half-height were computed. Latency was defined as the time from the touch onset to the first point exceeding half height (green circle). Shaded area, SEM. (C) Bootstrap procedure was performed to estimate the reliability of latency estimation. A narrow 90% CI indicates reliable estimation. (D-E) Same as B, C for a SOM neuron.

**Supplemental Figure 3.**
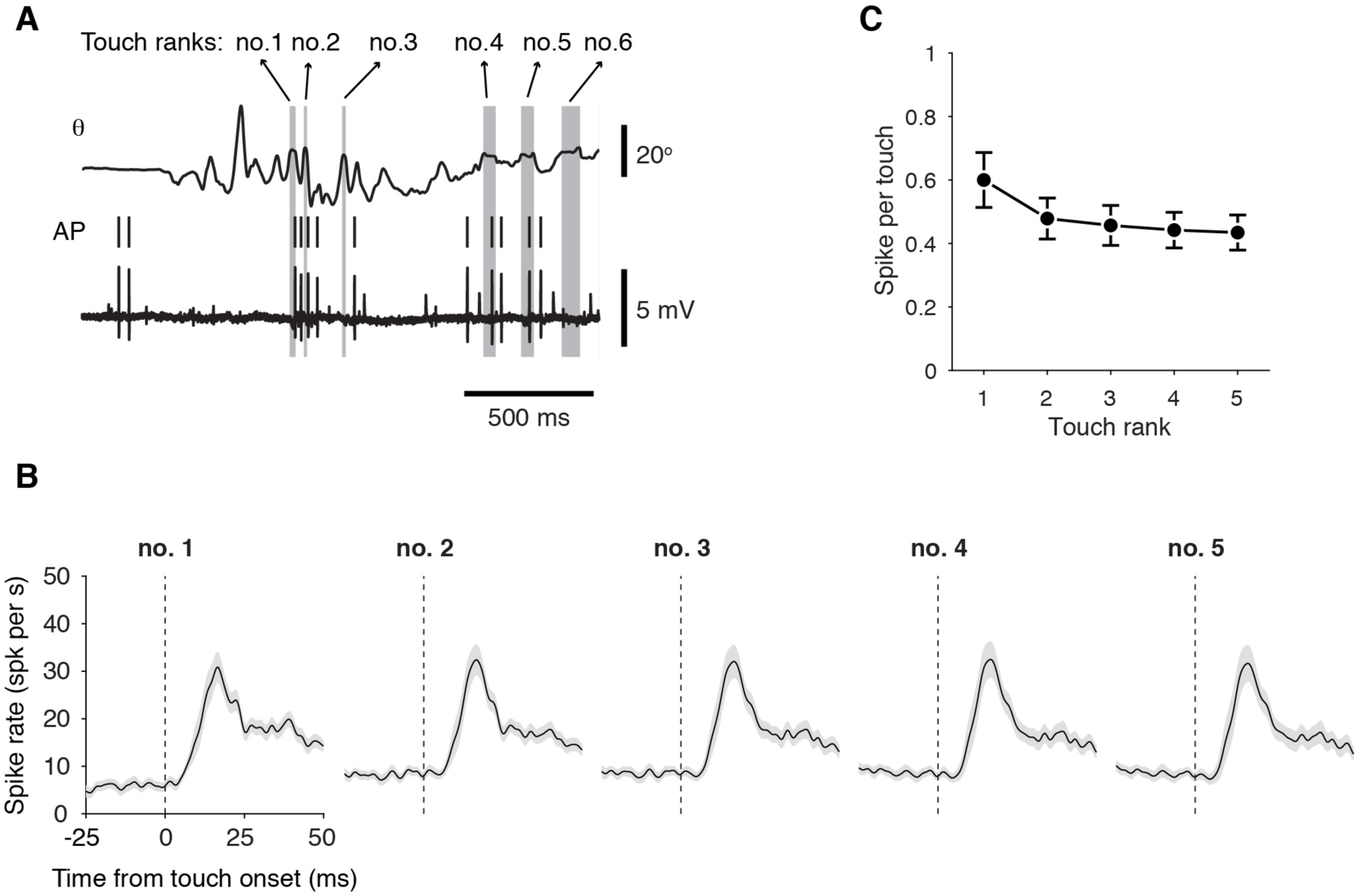
Touch-Evoked Spiking in SOM Neurons Does Not Require an Accumulation from Multiple Whisker Touches. (A) Definition of touch rank during a whisking bout. Touches within a whisking bout were typically 100 ms apart (median, 114 ms, 25^th^ to 75^th^ percentiles, 63 and 170 ms). (B) Touch-evoked responses (spike density function; Gaussian kernel, 1 ms) in SOM interneurons are grouped according to the order of touches during repetitive palpations (n = 58 neurons). Shaded areas, SEM. (C) Spikes per touch as a function of touch rank.

**Supplemental Figure 4.**
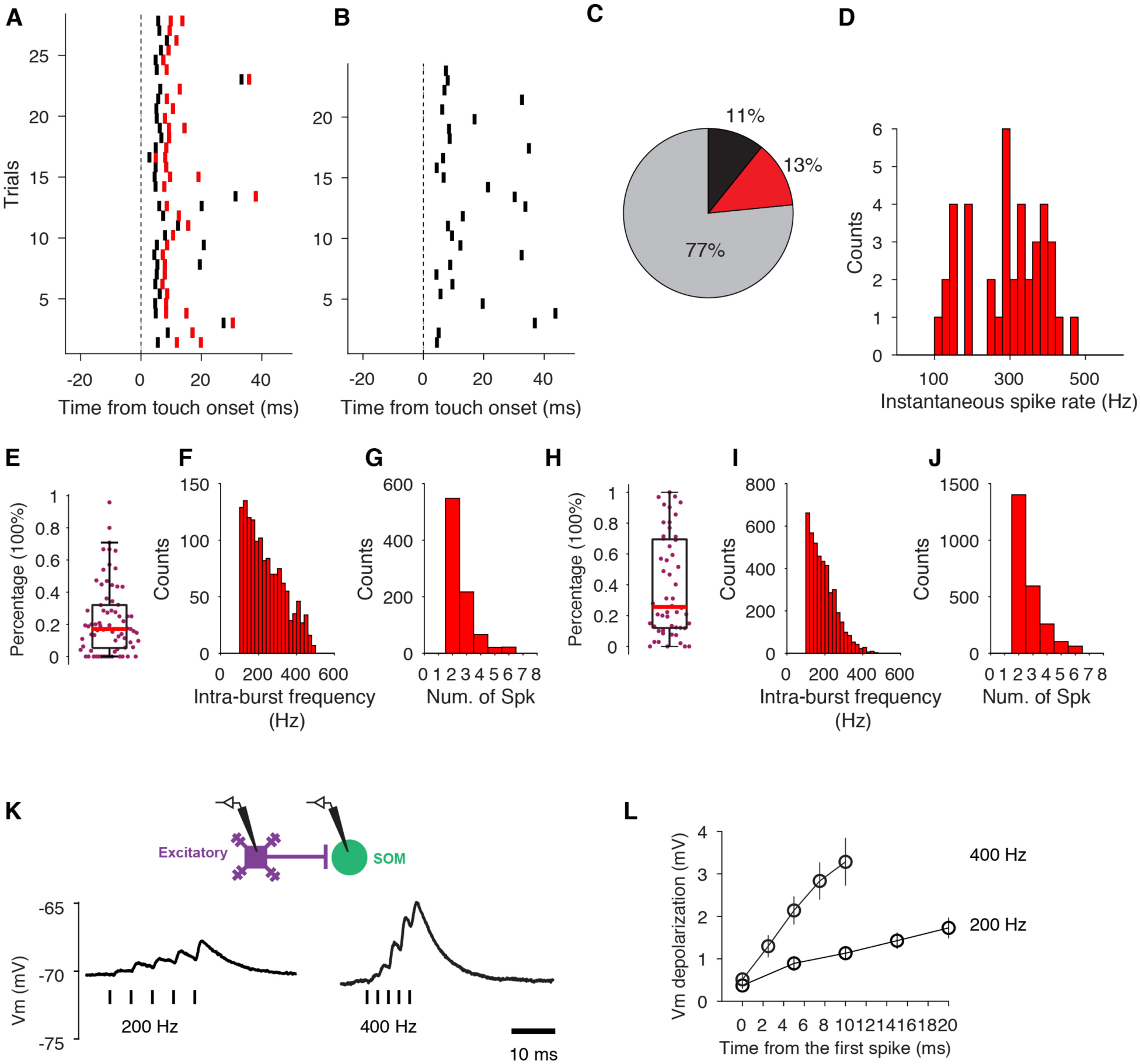
Burst Firing in E Neurons Could Enhance Depolarization in Postsynaptic SOM Neurons. (A) Touch triggered burst (inter-spike interval <10 ms) of spikes in an example L4 E neuron. Red, spikes with a preceding inter-spike interval of less than 10 ms. (B) Touch triggered non-burst firing in the same neuron. (C) Distribution of touch trials with no evoked spikes (gray), non-burst spikes (black), and burst spikes (red) in the example neuron. (D) Distribution of intra-burst spike frequency for all detected bursts in the example neuron. (E) Boxplot for the percentage of trials with evoked bursts over trials with evoked spiking (burst and non-burst) for all L4 E neurons. Red, median. (F) Distribution of intra-burst spike frequency for all bursts in all L4 E neurons (n = 73 cells). (G) Distribution of spike number in single bursts for all L4 E neurons. (H-J) Same as E-G for L5/6 E neurons (n = 53 cells). (K) Schematic, Paired recording of connected E and SOM neurons in brain slices. Bottom, example EPSP in an L4 SOM neuron evoked by high-frequency (200 or 400 Hz) spike train in a presynaptic E neuron (spikes indicated by vertical bars). Traces are trial-averaged responses. (L) Peak depolarization as a function of time during a 5-spike presynaptic spike train at 200 or 400 Hz (n =10 pairs).

**Supplemental Figure 5.**
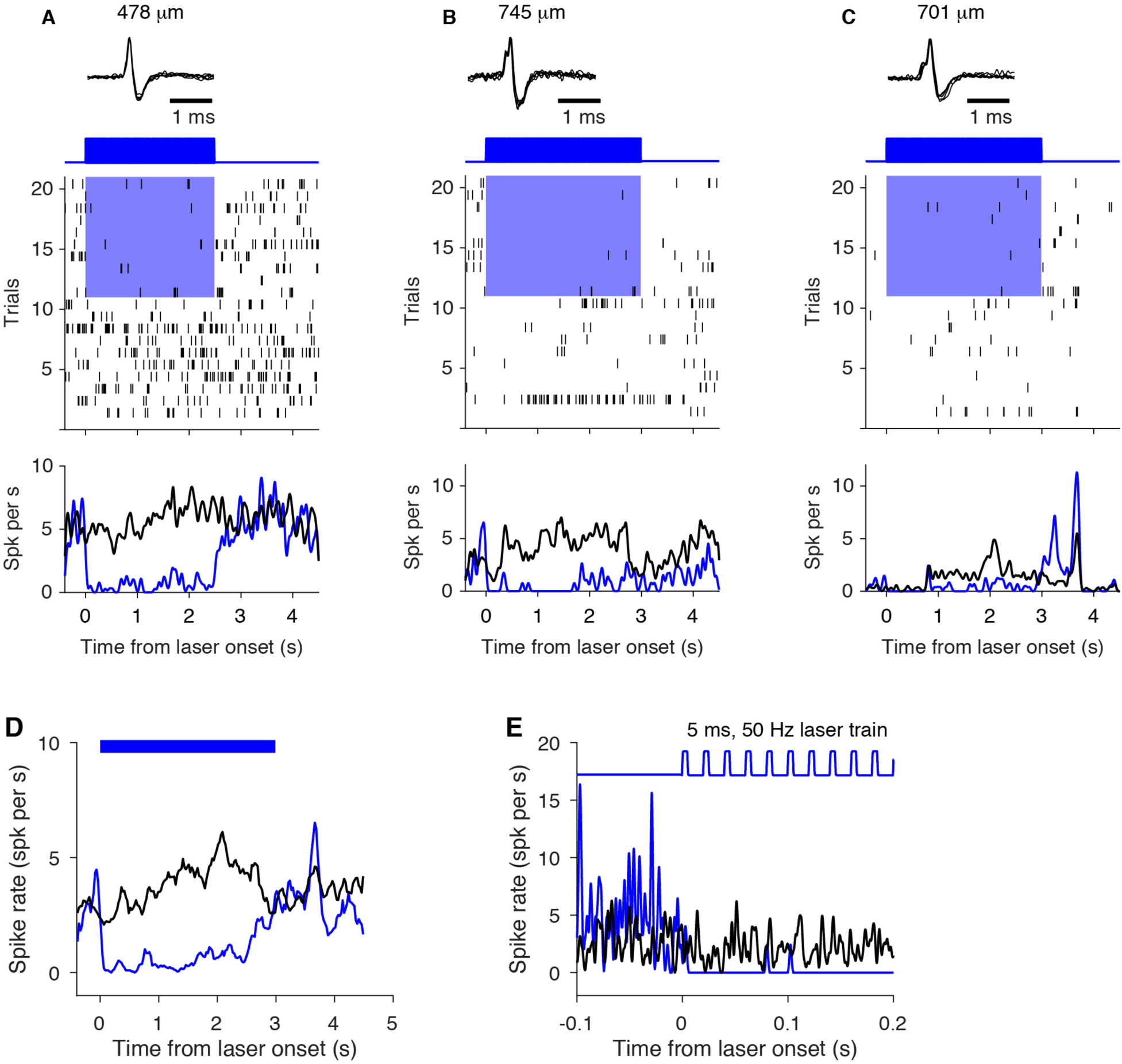
VIP Neuron Stimulation Inhibits Putative SOM Neurons. (A-C) Three putative SOM interneurons defined by their rapid suppression in response to VIP neuron stimulation. Their spike waveforms, spike raster plots, and PSTHs under control (black) and VIP stimulation (blue) conditions are shown. Note the characteristic spike rising phase (cf. Fig. S1) for cells in B and C. Recording depths (manipulator readings) are listed. (D) Averaged PSTHs of 3 putative SOM interneurons. Blue bars denote laser stimulation. (E) Averaged PSTH of 3 putative SOM interneurons at fine time scale around the laser onset.

## REFERENCES

Agmon, A., and Connors, B.W. (1991). Thalamocortical responses of mouse somatosensory (barrel) cortexin vitro. Neuroscience 41, 365–379.

Agmon, A., and Connors, B.W. (1992). Correlation between intrinsic firing patterns and thalamocortical synaptic responses of neurons in mouse barrel cortex. J. Neurosci. 12, 319–329.

Armstrong-James, M., Fox, K., and Das-Gupta, A. (1992). Flow of excitation within rat barrel cortex on striking a single vibrissa. J. Neurophysiol. 68, 1345–1358.

Audette, N.J., Urban-Ciecko, J., Matsushita, M., and Barth, A.L. (2018). POm Thalamocortical Input Drives Layer-Specific Microcircuits in Somatosensory Cortex. Cereb. Cortex N. Y. N 1991. 28, 1312–1328.

Beierlein, M., Gibson, J.R., and Connors, B.W. (2003). Two dynamically distinct inhibitory networks in layer 4 of the neocortex. J. Neurophysiol. 90, 2987–3000.

van Brederode, J.F., Helliesen, M.K., and Hendrickson, A.E. (1991). Distribution of the calcium-binding proteins parvalbumin and calbindin-D28k in the sensorimotor cortex of the rat. Neuroscience 44, 157–171.

Bruno, R.M., and Simons, D.J. (2002). Feedforward mechanisms of excitatory and inhibitory cortical receptive fields. J. Neurosci. Off. J. Soc. Neurosci. 22, 10966–10975.

Bureau, I., von Saint Paul, F., and Svoboda, K. (2006). Interdigitated paralemniscal and lemniscal pathways in the mouse barrel cortex. PLoS Biol. 4, e382.

Campagner, D., Evans, M.H., Bale, M.R., Erskine, A., and Petersen, R.S. (2016). Prediction of primary somatosensory neuron activity during active tactile exploration. ELife 5.

Chen, N., Sugihara, H., and Sur, M. (2015). An acetylcholine-activated microcircuit drives temporal dynamics of cortical activity. Nat. Neurosci. 18, 892–902.

Clack, N.G., O’Connor, D.H., Huber, D., Petreanu, L., Hires, A., Peron, S., Svoboda, K., and Myers, E.W. (2012). Automated tracking of whiskers in videos of head fixed rodents. PLoS Comput. Biol. 8, e1002591.

Constantinople, C.M., and Bruno, R.M. (2013). Deep Cortical Layers Are Activated Directly by Thalamus. Science 340, 1591–1594.

Crochet, S., Poulet, J.F.A., Kremer, Y., and Petersen, C.C.H. (2011). Synaptic mechanisms underlying sparse coding of active touch. Neuron 69, 1160–1175.

Cruikshank, S.J., Lewis, T.J., and Connors, B.W. (2007). Synaptic basis for intense thalamocortical activation of feedforward inhibitory cells in neocortex. Nat. Neurosci. 10, 462–468.

Cruikshank, S.J., Urabe, H., Nurmikko, A.V., and Connors, B.W. (2010). Pathway-specific feedforward circuits between thalamus and neocortex revealed by selective optical stimulation of axons. Neuron 65, 230–245.

Dantzker, J.L., and Callaway, E.M. (2000). Laminar sources of synaptic input to cortical inhibitory interneurons and pyramidal neurons. Nat. Neurosci. 3, 701–707.

Dávid, C., Schleicher, A., Zuschratter, W., and Staiger, J.F. (2007). The innervation of parvalbumin-containing interneurons by VIP-immunopositive interneurons in the primary somatosensory cortex of the adult rat. Eur. J. Neurosci. 25, 2329–2340.

Dipoppa, M., Ranson, A., Krumin, M., Pachitariu, M., Carandini, M., and Harris, K.D. (2018). Vision and Locomotion Shape the Interactions between Neuron Types in Mouse Visual Cortex. Neuron 98, 602–615.e8.

Feldmeyer, D., Qi, G., Emmenegger, V., and Staiger, J.F. (2018). Inhibitory interneurons and their circuit motifs in the many layers of the barrel cortex. Neuroscience 368, 132–151.

Fu, Y., Tucciarone, J.M., Espinosa, J.S., Sheng, N., Darcy, D.P., Nicoll, R.A., Huang, Z.J., and Stryker, M.P. (2014). A cortical circuit for gain control by behavioral state. Cell 156, 1139–1152.

Gabernet, L., Jadhav, S.P., Feldman, D.E., Carandini, M., and Scanziani, M. (2005). Somatosensory integration controlled by dynamic thalamocortical feed-forward inhibition. Neuron 48, 315–327.

Gentet, L.J., Avermann, M., Matyas, F., Staiger, J.F., and Petersen, C.C.H. (2010). Membrane potential dynamics of GABAergic neurons in the barrel cortex of behaving mice. Neuron 65, 422–435.

Gentet, L.J., Kremer, Y., Taniguchi, H., Huang, Z.J., Staiger, J.F., and Petersen, C.C.H. (2012). Unique functional properties of somatostatin-expressing GABAergic neurons in mouse barrel cortex. Nat. Neurosci. 15, 607–612.

Gibson, J.R., Beierlein, M., and Connors, B.W. (1999). Two networks of electrically coupled inhibitory neurons in neocortex. Nature 402, 75–79.

Groh, A., Kock, C.P.J. de, Wimmer, V.C., Sakmann, B., and Kuner, T. (2008). Driver or Coincidence Detector: Modal Switch of a Corticothalamic Giant Synapse Controlled by Spontaneous Activity and Short-Term Depression. J. Neurosci. 28, 9652–9663.

Guo, Z.V., Li, N., Huber, D., Ophir, E., Gutnisky, D., Ting, J.T., Feng, G., and Svoboda, K. (2014a). Flow of cortical activity underlying a tactile decision in mice. Neuron 81, 179–194.

Guo, Z.V., Hires, S.A., Li, N., O’Connor, D.H., Komiyama, T., Ophir, E., Huber, D., Bonardi, C., Morandell, K., Gutnisky, D., et al. (2014b). Procedures for behavioral experiments in head-fixed mice. PloS One 9, e88678.

Gutnisky, D.A., Yu, J., Hires, S.A., To, M.-S., Bale, M.R., Svoboda, K., and Golomb, D. (2017). Mechanisms underlying a thalamocortical transformation during active tactile sensation. PLoS Comput. Biol. 13, e1005576.

Hioki, H., Okamoto, S., Konno, M., Kameda, H., Sohn, J., Kuramoto, E., Fujiyama, F., and Kaneko, T. (2013). Cell type-specific inhibitory inputs to dendritic and somatic compartments of parvalbumin-expressing neocortical interneuron. J. Neurosci. Off. J. Soc. Neurosci. 33, 544–555.

Hires, S.A., Gutnisky, D.A., Yu, J., O’Connor, D.H., and Svoboda, K. (2015). Low-noise encoding of active touch by layer 4 in the somatosensory cortex. ELife 4.

Hong, Y.K., Lacefield, C.O., Rodgers, C.C., and Bruno, R.M. (2018). Sensation, movement and learning in the absence of barrel cortex. Nature 561, 542–546.

Hooks, B.M., Hires, S.A., Zhang, Y.-X., Huber, D., Petreanu, L., Svoboda, K., and Shepherd, G.M.G. (2011). Laminar analysis of excitatory local circuits in vibrissal motor and sensory cortical areas. PLoS Biol. 9, e1000572.

Hu, H., and Agmon, A. (2016). Differential Excitation of Distally versus Proximally Targeting Cortical Interneurons by Unitary Thalamocortical Bursts. J. Neurosci. Off. J. Soc. Neurosci. 36, 6906–6916.

Hu, H., Cavendish, J.Z., and Agmon, A. (2013). Not all that glitters is gold: off-target recombination in the somatostatin-IRES-Cre mouse line labels a subset of fast-spiking interneurons. Front. Neural Circuits 7, 195.

Jackson, J., Ayzenshtat, I., Karnani, M.M., and Yuste, R. (2016). VIP+ interneurons control neocortical activity across brain states. J. Neurophysiol. 115, 3008–3017.

Jiang, X., Shen, S., Cadwell, C.R., Berens, P., Sinz, F., Ecker, A.S., Patel, S., and Tolias, A.S. (2015). Principles of connectivity among morphologically defined cell types in adult neocortex. Science 350, aac9462–aac9462.

Kinnischtzke, A.K., Simons, D.J., and Fanselow, E.E. (2014). Motor cortex broadly engages excitatory and inhibitory neurons in somatosensory barrel cortex. Cereb. Cortex N. Y. N 1991 24, 2237–2248.

de Kock, C.P.J., Bruno, R.M., Spors, H., and Sakmann, B. (2007). Layer-and cell-type-specific suprathreshold stimulus representation in rat primary somatosensory cortex. J. Physiol. 581, 139–154.

Kvitsiani, D., Ranade, S., Hangya, B., Taniguchi, H., Huang, J.Z., and Kepecs, A. (2013). Distinct behavioural and network correlates of two interneuron types in prefrontal cortex. Nature 498, 363–366.

Lee, S., Hjerling-Leffler, J., Zagha, E., Fishell, G., and Rudy, B. (2010). The largest group of superficial neocortical GABAergic interneurons expresses ionotropic serotonin receptors. J. Neurosci. Off. J. Soc. Neurosci. 30, 16796–16808.

Lee, S., Kruglikov, I., Huang, Z.J., Fishell, G., and Rudy, B. (2013). A disinhibitory circuit mediates motor integration in the somatosensory cortex. Nat. Neurosci. 16, 1662–1670.

Lefort, S., Tomm, C., Floyd Sarria, J.-C., and Petersen, C.C.H. (2009). The excitatory neuronal network of the C2 barrel column in mouse primary somatosensory cortex. Neuron 61, 301–316.

Lima, S.Q., Hromádka, T., Znamenskiy, P., and Zador, A.M. (2009). PINP: A New Method of Tagging Neuronal Populations for Identification during In Vivo Electrophysiological Recording. PLOS ONE 4, e6099.

Lu, S.-M., and Lin, R.C.-S. (1993). Thalamic Afferents of the Rat Barrel Cortex: A Light-and Electron-Microscopic Study Using Phaseolus vulgaris Leucoagglutinin as an Anterograde Tracer. Somatosens. Mot. Res. 10, 1–16.

Ma, Y., Hu, H., Berrebi, A.S., Mathers, P.H., and Agmon, A. (2006). Distinct Subtypes of Somatostatin-Containing Neocortical Interneurons Revealed in Transgenic Mice. J. Neurosci. 26, 5069–5082.

Ma, Y., Hu, H., and Agmon, A. (2012). Short-Term Plasticity of Unitary Inhibitory-to-Inhibitory Synapses Depends on the Presynaptic Interneuron Subtype. J. Neurosci. 32, 983–988.

Madisen, L., Zwingman, T.A., Sunkin, S.M., Oh, S.W., Zariwala, H.A., Gu, H., Ng, L.L., Palmiter, R.D., Hawrylycz, M.J., Jones, A.R., et al. (2010). A robust and high-throughput Cre reporting and characterization system for the whole mouse brain. Nat. Neurosci. 13, 133–140.

Madisen, L., Mao, T., Koch, H., Zhuo, J., Berenyi, A., Fujisawa, S., Hsu, Y.-W.A., Garcia, A.J., Gu, X., Zanella, S., et al. (2012). A toolbox of Cre-dependent optogenetic transgenic mice for light-induced activation and silencing. Nat. Neurosci. 15, 793–802.

Mao, T., Kusefoglu, D., Hooks, B.M., Huber, D., Petreanu, L., and Svoboda, K. (2011). Long-Range Neuronal Circuits Underlying the Interaction between Sensory and Motor Cortex. Neuron 72, 111–123.

Masri, R., Bezdudnaya, T., Trageser, J.C., and Keller, A. (2008). Encoding of stimulus frequency and sensor motion in the posterior medial thalamic nucleus. J. Neurophysiol. 100, 681–689.

Moore, A.K., and Wehr, M. (2013). Parvalbumin-expressing inhibitory interneurons in auditory cortex are well-tuned for frequency. J. Neurosci. Off. J. Soc. Neurosci. 33, 13713–13723.

Moore, J.D., Mercer Lindsay, N., Deschênes, M., and Kleinfeld, D. (2015). Vibrissa Self-Motion and Touch Are Reliably Encoded along the Same Somatosensory Pathway from Brainstem through Thalamus. PLoS Biol. 13, e1002253.

Muñoz, W., Tremblay, R., and Rudy, B. (2014). Channelrhodopsin-assisted patching: in vivo recording of genetically and morphologically identified neurons throughout the brain. Cell Rep. 9, 2304–2316.

Muñoz, W., Tremblay, R., Levenstein, D., and Rudy, B. (2017). Layer-specific modulation of neocortical dendritic inhibition during active wakefulness. Science 355, 954–959.

Nawrot, M., Aertsen, A., and Rotter, S. (1999). Single-trial estimation of neuronal firing rates: From single-neuron spike trains to population activity. J. Neurosci. Methods 94, 81–92.

O’Connor, D.H., Clack, N.G., Huber, D., Komiyama, T., Myers, E.W., and Svoboda, K. (2010a). Vibrissa-based object localization in head-fixed mice. J. Neurosci. Off. J. Soc. Neurosci. 30, 1947–1967.

O’Connor, D.H., Peron, S.P., Huber, D., and Svoboda, K. (2010b). Neural activity in barrel cortex underlying vibrissa-based object localization in mice. Neuron 67, 1048–1061.

O’Connor, D.H., Hires, S.A., Guo, Z.V., Li, N., Yu, J., Sun, Q.-Q., Huber, D., and Svoboda, K. (2013). Neural coding during active somatosensation revealed using illusory touch. Nat. Neurosci. 16, 958–965.

Pakan, J.M., Lowe, S.C., Dylda, E., Keemink, S.W., Currie, S.P., Coutts, C.A., and Rochefort, N.L. (2016). Behavioral-state modulation of inhibition is context-dependent and cell type specific in mouse visual cortex. ELife 5.

Pammer, L., O’Connor, D.H., Hires, S.A., Clack, N.G., Huber, D., Myers, E.W., and Svoboda, K. (2013). The mechanical variables underlying object localization along the axis of the whisker. J. Neurosci. Off. J. Soc. Neurosci. 33, 6726–6741.

Petersen, R.S., Brambilla, M., Bale, M.R., Alenda, A., Panzeri, S., Montemurro, M.A., and Maravall, M. (2008). Diverse and Temporally Precise Kinetic Feature Selectivity in the VPm Thalamic Nucleus. Neuron 60, 890–903.

Petreanu, L., Mao, T., Sternson, S.M., and Svoboda, K. (2009). The subcellular organization of neocortical excitatory connections. Nature 457, 1142–1145.

Pfeffer, C.K., Xue, M., He, M., Huang, Z.J., and Scanziani, M. (2013). Inhibition of inhibition in visual cortex: the logic of connections between molecularly distinct interneurons. Nat. Neurosci. 16, 1068–1076.

Pi, H.-J., Hangya, B., Kvitsiani, D., Sanders, J.I., Huang, Z.J., and Kepecs, A. (2013). Cortical interneurons that specialize in disinhibitory control. Nature 503, 521–524.

Pinault, D. (1996). A novel single-cell staining procedure performed in vivo under electrophysiological control: morpho-functional features of juxtacellularly labeled thalamic cells and other central neurons with biocytin or Neurobiotin. J. Neurosci. Methods 65, 113–136.

Pinault, D. (2005). A new stabilizing craniotomy–duratomy technique for single-cell anatomo-electrophysiological exploration of living intact brain networks. J. Neurosci. Methods 141, 231–242.

Porter, J.T., Johnson, C.K., and Agmon, A. (2001). Diverse Types of Interneurons Generate Thalamus-Evoked Feedforward Inhibition in the Mouse Barrel Cortex. J. Neurosci. 21, 2699–2710.

Poulet, J.F.A., Fernandez, L.M.J., Crochet, S., and Petersen, C.C.H. (2012). Thalamic control of cortical states. Nat. Neurosci. 15, 370–372.

Prönneke, A., Scheuer, B., Wagener, R.J., Möck, M., Witte, M., and Staiger, J.F. (2015). Characterizing VIP Neurons in the Barrel Cortex of VIPcre/tdTomato Mice Reveals Layer-Specific Differences. Cereb. Cortex 25, 4854–4868.

Reyes, A., Lujan, R., Rozov, A., Burnashev, N., Somogyi, P., and Sakmann, B. (1998). Target-cell-specific facilitation and depression in neocortical circuits. Nat. Neurosci. 1, 279–285.

Rudy, B., Fishell, G., Lee, S., and Hjerling-Leffler, J. (2011). Three groups of interneurons account for nearly 100% of neocortical GABAergic neurons. Dev. Neurobiol. 71, 45–61.

Sachidhanandam, S., Sermet, B.S., and Petersen, C.C.H. (2016). Parvalbumin-Expressing GABAergic Neurons in Mouse Barrel Cortex Contribute to Gating a Goal-Directed Sensorimotor Transformation. Cell Rep. 15, 700–706.

Severson, K.S., Xu, D., Van de Loo, M., Bai, L., Ginty, D.D., and O’Connor, D.H. (2017). Active Touch and Self-Motion Encoding by Merkel Cell-Associated Afferents. Neuron 94, 666–676.e9.

Shepherd, G.M.G., and Svoboda, K. (2005). Laminar and columnar organization of ascending excitatory projections to layer 2/3 pyramidal neurons in rat barrel cortex. J. Neurosci. Off. J. Soc. Neurosci. 25, 5670–5679.

Simons, D.J., and Carvell, G.E. (1989). Thalamocortical response transformation in the rat vibrissa/barrel system. J. Neurophysiol. 61, 311–330.

Sohn, J., Okamoto, S., Kataoka, N., Kaneko, T., Nakamura, K., and Hioki, H. (2016). Differential Inputs to the Perisomatic and Distal-Dendritic Compartments of VIP-Positive Neurons in Layer 2/3 of the Mouse Barrel Cortex. Front. Neuroanat. 10.

Staiger, J.F., Zilles, K., and Freund, T.F. (1996). Innervation of VIP-immunoreactive neurons by the ventroposteromedial thalamic nucleus in the barrel cortex of the rat. J. Comp. Neurol. 367, 194–204.

Staiger, J.F., Freund, T.F., and Zilles, K. (1997). Interneurons Immunoreactive for Vasoactive Intestinal Polypeptide (VIP) are Extensively Innervated by Parvalbumin-Containing Boutons in Rat Primary Somatosensory Cortex. Eur. J. Neurosci. 9, 2259–2268.

Staiger, J.F., Schubert, D., Zuschratter, W., Kötter, R., and Luhmann, H.J. (2002). Innervation of interneurons immunoreactive for VIP by intrinsically bursting pyramidal cells and fast-spiking interneurons in infragranular layers of juvenile rat neocortex. Eur. J. Neurosci. 16, 11–20.

Sun, Q.-Q., Huguenard, J.R., and Prince, D.A. (2006). Barrel cortex microcircuits: thalamocortical feedforward inhibition in spiny stellate cells is mediated by a small number of fast-spiking interneurons. J. Neurosci. Off. J. Soc. Neurosci. 26, 1219–1230.

Suter, B.A., O’Connor, T., Iyer, V., Petreanu, L., Hooks, B.M., Kiritani, T., Svoboda, K., and Shepherd, G.M.G. (2010). Ephus: Multipurpose Data Acquisition Software for Neuroscience Experiments. Front. Neural Circuits 4.

Swadlow, H.A. (1995). Influence of VPM afferents on putative inhibitory interneurons in S1 of the awake rabbit: evidence from cross-correlation, microstimulation, and latencies to peripheral sensory stimulation. J. Neurophysiol. 73, 1584–1599.

Taniguchi, H., He, M., Wu, P., Kim, S., Paik, R., Sugino, K., Kvitsani, D., Fu, Y., Lu, J., Lin, Y., et al. (2011). A Resource of Cre Driver Lines for Genetic Targeting of GABAergic Neurons in Cerebral Cortex. Neuron 71, 995–1013.

Tasic, B., Yao, Z., Graybuck, L.T., Smith, K.A., Nguyen, T.N., Bertagnolli, D., Goldy, J., Garren, E., Economo, M.N., Viswanathan, S., et al. (2018). Shared and distinct transcriptomic cell types across neocortical areas. Nature 563, 72–78.

Tremblay, R., Lee, S., and Rudy, B. (2016). GABAergic Interneurons in the Neocortex: From Cellular Properties to Circuits. Neuron 91, 260–292.

Urbain, N., Salin, P.A., Libourel, P.-A., Comte, J.-C., Gentet, L.J., and Petersen, C.C.H. (2015). Whisking-Related Changes in Neuronal Firing and Membrane Potential Dynamics in the Somatosensory Thalamus of Awake Mice. Cell Rep. 13, 647–656.

Wall, N.R., De La Parra, M., Sorokin, J.M., Taniguchi, H., Huang, Z.J., and Callaway, E.M. (2016). Brain-Wide Maps of Synaptic Input to Cortical Interneurons. J. Neurosci. Off. J. Soc. Neurosci. 36, 4000–4009.

Wimmer, V.C., Bruno, R.M., de Kock, C.P.J., Kuner, T., and Sakmann, B. (2010). Dimensions of a projection column and architecture of VPM and POm axons in rat vibrissal cortex. Cereb. Cortex N. Y. N 1991 20, 2265–2276.

Xu, H., Jeong, H.-Y., Tremblay, R., and Rudy, B. (2013). Neocortical Somatostatin-expressing GABAergic Interneurons Disinhibit the Thalamorecipient Layer 4. Neuron 77, 155–167.

Yu, J., Gutnisky, D.A., Hires, S.A., and Svoboda, K. (2016). Layer 4 fast-spiking interneurons filter thalamocortical signals during active somatosensation. Nat. Neurosci. 19, 1647–1657.

